# Transcriptomic-based quantification of the epithelial-hybrid-mesenchymal spectrum across biological contexts

**DOI:** 10.1101/2021.09.09.458982

**Authors:** Susmita Mandal, Tanishq Tejaswi, Rohini Janivara, Syamanthak Srikrishnan, Pradipti Thakur, Sarthak Sahoo, Priyanka Chakraborty, Sukhwinder Singh Sohal, Herbert Levine, Jason T George, Mohit Kumar Jolly

**Author notes:** To whom correspondence should be addressed (J.T.G.), (M.K.J.).

## Abstract

Epithelial-mesenchymal plasticity (EMP) underlies embryonic development, wound healing, and cancer metastasis and fibrosis. Cancer cells exhibiting EMP often have more aggressive behavior, characterized by drug resistance, and tumor-initiating and immuno-evasive traits. Thus, the EMP status of cancer cells can be a critical indicator of patient prognosis. Here, we compare three distinct transcriptomic-based metrics – each derived using a different gene list and algorithm – that quantify the EMP spectrum. Our results for 96 cancer-related RNA-seq datasets reveal a high degree of concordance among these metrics in quantifying the extent of EMP. Moreover, each metric, despite being trained on cancer expression profiles, recapitulates the expected changes in EMP scores for non-cancer contexts such as lung fibrosis and cellular reprogramming into induced pluripotent stem cells. Thus, we offer a scoring platform to quantify the extent of EMP *in vitro* and *in vivo* for diverse biological applications including cancer.

## Introduction

Epithelial-Mesenchymal Plasticity (EMP) is an important feature of cancer metastasis and therapy resistance, the two major clinical challenges that claim the majority of cancer-related deaths (Gupta and Massague, 2006). EMP involves dynamic and reversible switching among multiple phenotypes along the epithelial-hybrid-mesenchymal spectrum. It encompasses both EMT (Epithelial-to-Mesenchymal Transition) and MET (Mesenchymal-to-Epithelial Transition). Originally considered to be binary transitions, EMT and MET are both now understood as multistep processes, and cells can execute these programs to varying degrees, enabling one or more hybrid epithelial/mesenchymal (E/M) phenotypes (Jolly and Levine, 2017; Nieto et al., 2016; Pal et al., 2021). EMP usually entails changes in cell-cell adhesion, migration, and invasion; while EMT is often involved with cells escaping the primary tumor and initiating metastasis, MET is thought to be important for colonization, the last step of metastasis. Besides these features, EMP is also implicated in conferring tumor-initiation potential (Morel et al., 2008; Pasani et al., 2021), immune evasion (Chen et al., 2014; Dongre et al., 2017; S. C. Tripathi et al., 2016), and resistance to various chemotherapeutic drugs and targeted therapies (Creighton et al., 2009; Sahoo et al., 2021a; Wang et al., 2009). Thus, EMP can be considered as the “motor of cellular plasticity” (Brabletz and Brabletz, 2010), which enhances cancer cell fitness in a variety of biological contexts.

Recent preclinical and clinical observations have suggested high metastatic potential of hybrid E/M phenotypes, and their association with worse patient survival across cancer types (Bierie et al., 2017; Godin et al., 2020; Huang et al., 2013; Kröger et al., 2019; Pastushenko and Blanpain, 2019; Puram et al., 2017; Sahoo et al., 2021b; Simeonov et al., 2021). Hybrid E/M phenotypes have also been observed in circulating tumor cells (CTCs); their higher frequency is often concomitant with worse clinicopathological features (Bocci et al., 2021; Lecharpentier et al., 2011; Saxena et al., 2019; Yu et al., 2013). The ability of hybrid E/M cells to form clusters of CTCs can also escalate their metastatic fitness, given the disproportionately high metastatic burden of CTC clusters (Aceto et al., 2014; Cheung et al., 2016; Jolly et al., 2017). Given the context-specific diversity of hybrid E/M phenotypes (Jolly et al., 2021), it is imperative that EMP be quantified as a continuum or spectrum, through integrating various experimental and/or computational methods.

At the transcriptomic level, various computational methods have been proposed to quantify the EMP spectrum by calculating the extent to which a given sample has undergone EMT/MET (hereafter, referred to as the ‘EMT score’). First, using gene expression from non-small cell lung cancer (NSCLC) cell lines and patients, a 76-gene EMT signature was identified and then used to derive a score (hereafter, referred to as the ‘76GS score’) based on the relative enrichment of expression levels of epithelial-associated genes. Thus, the higher the 76GS score, the more epithelial a given sample is (Byers et al., 2013; Guo et al., 2019). Second, a two-sample Kolmogorov-Smirnov method was used to calculate a score (hereafter, referred to as the ‘KS score’) on the interval [-1, +1] to depict the EMP status of cell lines and tumors. The higher the KS score, the more mesenchymal a sample is (Tan et al., 2014). Third, a multinomial logistic regression method implemented on NCI-60 expression data quantified the extent of EMT on the interval [0, 2] (hereafter, referred to as the ‘MLR score’) by calculating the probabilities for a given sample to belong to E, M or hybrid E/M states (George et al., 2017). Higher MLR scores depict a relatively enriched mesenchymal phenotype. A previous study compared these three methods – each of which utilizes a distinct gene list and algorithm – and observed that these methods were largely well-correlated with one another in terms of quantifying EMP across multiple microarray datasets (Chakraborty et al., 2020). This analysis suggested that 76GS scores correlated negatively with their MLR and KS counterparts, both of which positively correlated with one another. However, two key questions remain unanswered: a) can all three of these scoring metrics quantify the EMP spectrum for bulk and single-cell RNA-seq data with the same level of consistency? b) can these scores, all constructed on cancer-related datasets, be helpful in estimating the extent of EMP in non-cancer scenarios as well?

Here, we have addressed these limitations by analyzing multiple bulk and single-cell RNA-seq datasets, as well as investigating both microarray and RNA-seq datasets for two non-cancer cases where EMP has been reported: a) lung diseases - chronic obstructive pulmonary disease (COPD) and fibrosis (Jolly et al., 2018; Sohal, 2017) and b) reprogramming into induced pluripotent stem cells (iPSCs) (Lai et al., 2020). We demonstrate consistency amongst the EMT scoring metrics in quantifying the EMP spectrum across these biological contexts, as well as heterogeneity of EMP phenotypes in single-cell RNA-seq datasets. Finally, through a pan-cancer analysis of RNA-seq data available via The Cancer Genome Atlas (TCGA), we show that the association of EMP with patient survival is context-specific. Despite using diverse gene-sets and methodology to quantify EMP, a convergence of these three methods suggests possible commonalities in the different trajectories that cells undergoing EMT/MET can take in a high-dimensional landscape. Moreover, our results offer proof-of-principle that these metrics, all of which were derived based on cancer cells, can successfully quantify EMP in other useful non-cancer biological contexts too.

## Results

### EMT scoring methods show concordant trends across bulk RNA-seq datasets

We used the three different EMT scoring methods – 76GS, KS, and MLR – to quantify the extent of EMP in multiple RNA-seq datasets, as was previously done for microarray data (Chakraborty et al., 2020). Each method utilizes a distinct gene signature and underlying algorithm to compute an EMT score. The 76GS score is a weighted sum of the expression of 76 genes, where the weight factor is the correlation coefficient of that gene with the expression levels of CDH1 (E-cadherin), a canonical epithelial marker. Thus, a higher 76GS score corresponds to a more epithelial sample (Byers et al., 2013; Guo et al., 2019). The KS scoring method compares the empirical distribution function to the cumulative distribution function for epithelial and mesenchymal signatures identified in cell lines and tumors. The KS score is constructed by taking the maximal difference in these distributions for each predictor, followed by normalization by the number of predictors, thus taking values between −1 and +1. Positive (resp. negative) KS scores correspond to a relative enrichment of the mesenchymal (resp. epithelial) signature (Tan et al., 2014). The Multinomial Logistic Regression-based (MLR) method quantifies the extent of EMT on a scale of 0-2. MLR scores are calculated based on the probability of a given sample being assigned to the E, E/M, and M phenotypes. Thus, the higher the score, the more mesenchymal the sample is (George et al., 2017). While KS and 76GS methods operate on gene lists and can therefore be directly applied to both microarray and RNA-seq data, the MLR method utilizes the NCI-60 microarray data as training set for regression. Therefore, applying these methods for analyzing RNA-seq data needs further customization.

We extended the previous MLR framework trained on microarray-based transcriptomics of NCI-60 series to impute log_2_-normalized FPKM or TPM RNA-seq data. To achieve this, the log_2_ RNA-seq values for 3 predictors (CLDN7, VIM, CDH1) and 20 normalizers were linearly mapped to their corresponding microarray values (**Fig 1A**). This mapping was estimated for both FPKM- and TPM-normalized data by averaging over 24 previously published samples (Zhao et al., 2014) where log_2_ microarray and log_2_ RNA-seq expression signatures were simultaneously available (**Fig S1, S2**). The output of the updated MLR approach assigns a numerical EMT score, *S*, on the scale of [0, 2] based on the probability of a sample’s categorization into one of three groups: E, E/M and M.

**Figure 1.**
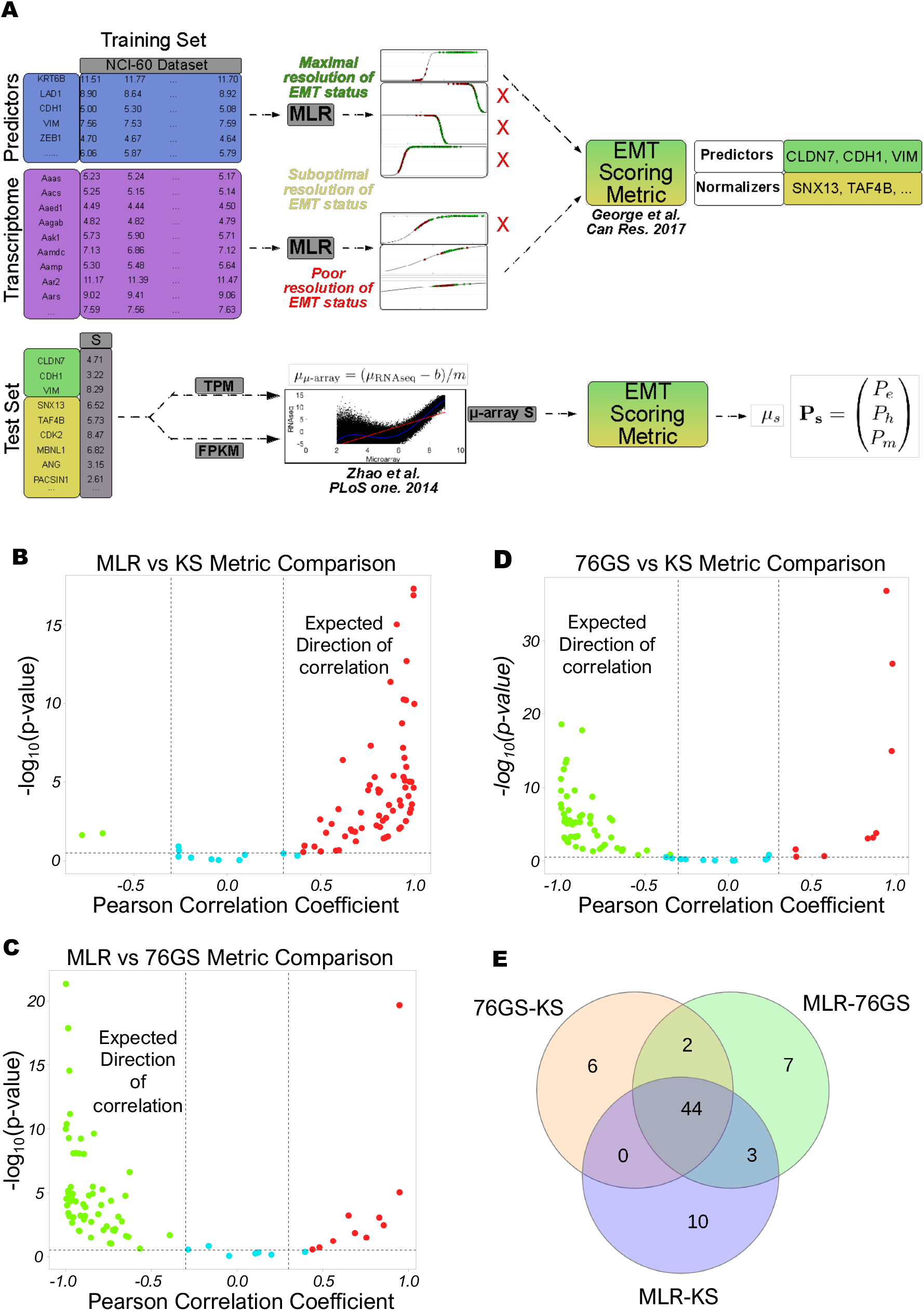
Schematic of the interconversion of MLR microarray and MLR RNA-seq scores and pairwise correlation plots of the three EMT scoring metrics. **A)** Adaptation of microarray-based MLR approach to predict RNA-seq-based input. Predictors (resp. normalizers) in the NCI-60 microarray training set were selected as done previously, based on their ability (resp. inability) to resolve the EMT status of the training set. Linear regression between RNA-seq and microarray data given in Zhao et al. 2014 was used to relate a unique value of RNA-seq signal to the microarray-based model, done for both FPKM and TPM values. **B-D**) Plots showing the pairwise correlation of EMT scoring metrics across 77 bulk RNA-Seq datasets, and for each sample –log_10_(p-value) is plotted as a function of Pearson’s correlation coefficient. Thresholds considered for correlation (R < −0.3 or R > 0.3; vertical dashed grey lines) and p-values (p < 0.05; horizontal dashed grey lines) are denoted. **E)** Venn diagram depicting the overlap of pairwise and full concordance for datasets that are significantly correlated in the expected direction.

To check the concordance of these three EMT scoring metrics, we calculated the 76GS, KS and MLR scores for 77 bulk high-throughput transcriptomics datasets. As expected, we found 76GS scores to be negatively correlated (r < −0.3; p < 0.045) with the MLR and KS scores, and found a positive correlation (r > 0.3; p < 0.05) between the MLR and KS scores across most datasets that contain cell lines and primary tumors across cancer types (**Fig 1B-D; Table S1**). 44 out of 77 (57.14%) datasets showed all three trends significantly (KS vs. MLR, MLR vs. 76GS and 76GS vs. KS). Additionally, 52 (67.53%) cases exhibit expected trends for 76GS vs. KS, compared to 56 (72.72%) for MLR vs. 76GS, and 57 (74.02%) for MLR vs. KS (**Fig 1E**). Thus, the MLR, 76GS, and KS scoring metrics show strong concordance among themselves for these 77 datasets.

Next, we investigated several individual datasets where EMT/MET phenomenon was induced in different tissues and cell lines. We found that the Py2T murine epithelial tumor cells that exhibit reversible EMT upon treatment with TGF-β *in vitro* had lower KS and MLR scores but higher 76GS scores as compared to MTΔEcad cells that represent irreversible EMT murine mammary gland tumor cells (**Fig 2A**; GSE118612) (Ishay-Ronen et al., 2019). Thus, all three EMT scores captured the expected trend of Py2T cells being more epithelial relative to MTΔEcad (murine breast cancer cells with ablated E-cadherin). Further, in mammary epithelial cells (MCF10A), depletion of Runx1 results in striking morphological changes consistent with EMT (Hong et al., 2017). Consistently, Runx1 depleted MCF10A cells had higher KS and MLR scores, but lower 76GS scores (**Fig 2B**; GSE85857). Similarly, TGF-β treated primary airway epithelial cells as well as TGF-β and EGF treated HeLa cells had a more mesenchymal profile as assessed by 76GS, KS and MLR scores, consistent with their reported experimental trends (**Fig 2C-D**; GSE72419, GSE61220) (Tian et al., 2015; V. Tripathi et al., 2016). These scores were also able to recapitulate *in vitro* observations that, while TGF-β treatment was able to induce EMT in MCF10A cells, the extent of EMT induced was decreased upon knockdown of ZEB1 (**Fig 2E**, GSE1248423) (Watanabe et al., 2019), a key EMT-inducing transcription factor in many cancers (Drápela et al., 2020). ZEB1 forms a mutually inhibitory feedback loop with GRHL2, a crucial MET-inducing factor, and knockdown of GRHL2 is known to push epithelial or hybrid E/M cells into a more mesenchymal phenotype (Chung et al., 2016; Cieply et al., 2013; Jolly et al., 2016; Mooney et al., 2017). Therefore, OVCA429 cells with GRHL2 knockdown had higher MLR and KS scores, but reduced 76GS scores, as compared to control, reflective of their more mesenchymal status (**Fig 2F**, GSE118407) (Chung et al., 2019). Similarly, *Grhl2*-null embryos had reduced levels of other gatekeepers of epithelial phenotype (*Ovol1, Ovol2* and miR-200 family (Aue et al., 2015; Chung et al., 2016; Jia et al., 2015)) and elevated levels of *Zeb1*, commensurate with their altered KS, 76GS and MLR scores (**Fig 2G**, GSE106130) (Carpinelli et al., 2020). ZEB1 is directly activated by Twist (Dave et al., 2011), another well-characterized EMT inducer (Yang et al., 2004). Thus, activation of Twist in HMLE (human mammary gland epithelial cells) corresponded to higher KS and MLR scores and reduced 76GS scores (**Fig 2I**; GSE139074).

**Figure 2.**
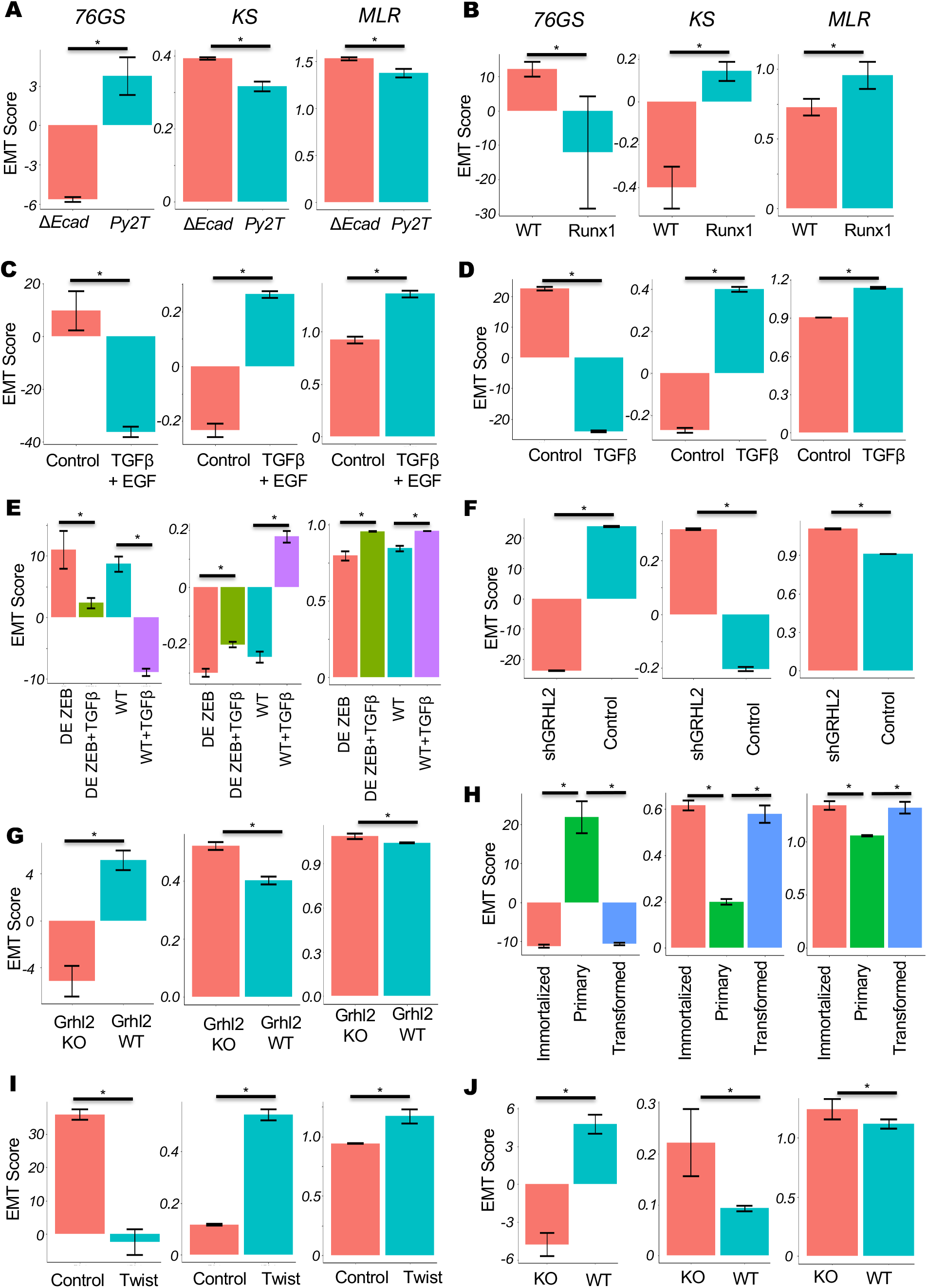
KS, 76GS and MLR scores for different EMT associated datasets. **A)** Py2T long term cells and mesenchymal breast cancer cells (MTΔECad) treated with different inhibitors (GSE118612). **B)** MCF10A cells with or without depletion of Runx1 (GSE85857). **C)** EMT induction in Hela cells by TGF-β + EGF (GSE72419) treatment. **D)** EMT induction in small airway epithelial cells by TGF-β (GSE61220). **E)** MCF10A cells with/without ZEB1 knocked out (KO) untreated or treated with TGF-β (GSE124843). **F)** OVCA4209 cells with/without GRHL2 knockdown (GSE118407). **G)** Grhl2 KO and wild type (WT) mice (GSE106130). **H)** Primary, pre-malignant immortalized, and Ras-transformed human mammary epithelial cells (GSE110677). **I)** EMT induction in HMLE/Twist-ER cells by tamoxifen (GSE139074). **J)** EMT status in HNF-1β-deficient mIMCD3 cell lines (GSE97770).

Similarly, these scoring metrics could recapitulate activation of EMT in pre-malignant immortalized and Ras-transformed HMECs (human mammary epithelial cells) as compared to primary HMECs (GSE110677; **Fig 2H**). Finally, in the context of renal fibrosis caused by loss of HNF-1β (Chan et al., 2018), HNF-1β deficient renal epithelial cells mIMCD3 showed upregulated mesenchymal traits relative to wild-type cells, as again captured by KS, 76GS and MLR scores (**Fig 2J**; GSE97770) (**Table S2**). Together, these case studies demonstrate that each scoring metric can capture the extent of EMT induced upon various perturbations, consistent with enrichment of EMT depicted by the Hallmark EMT geneset reported in MSigDB (Molecular Signature Database) (**Fig S3**) (Liberzon et al., 2011).

### Single-cell RNA-seq data analysis reveals heterogeneity along the EMP spectrum

After investigating bulk RNA-seq datasets, we calculated EMT scores for 17 single-cell RNA-seq datasets using 76GS, KS and MLR metrics. For example, in a dataset containing 5902 single cells isolated from 18 patients with oral cavity tumors (head and neck squamous cell carcinoma), we observed a negative correlation between 76GS and KS scores, and between 76GS and MLR ones, with however a positive correlation between the KS and MLR scores (**Fig 3A**; GSE103322) (Puram et al., 2017). This trend was largely seen across other single-cell RNA-seq datasets as well, where, like our previous results for bulk RNA-seq datasets, roughly 65% (11/17) of datasets showed negative correlation for 76GS vs. KS scores, 59% (10/17) datasets had negative correlation for MLR vs. 76GS, and 53% (9/17) exhibited a positive correlation for MLR vs. KS (**Fig 3B-C, S4A**). Thus, the concordant trends observed for these metrics using bulk transcriptomics were found to be conserved for single-cell RNA-seq datasets as well (**Table S3**).

**Figure 3.**
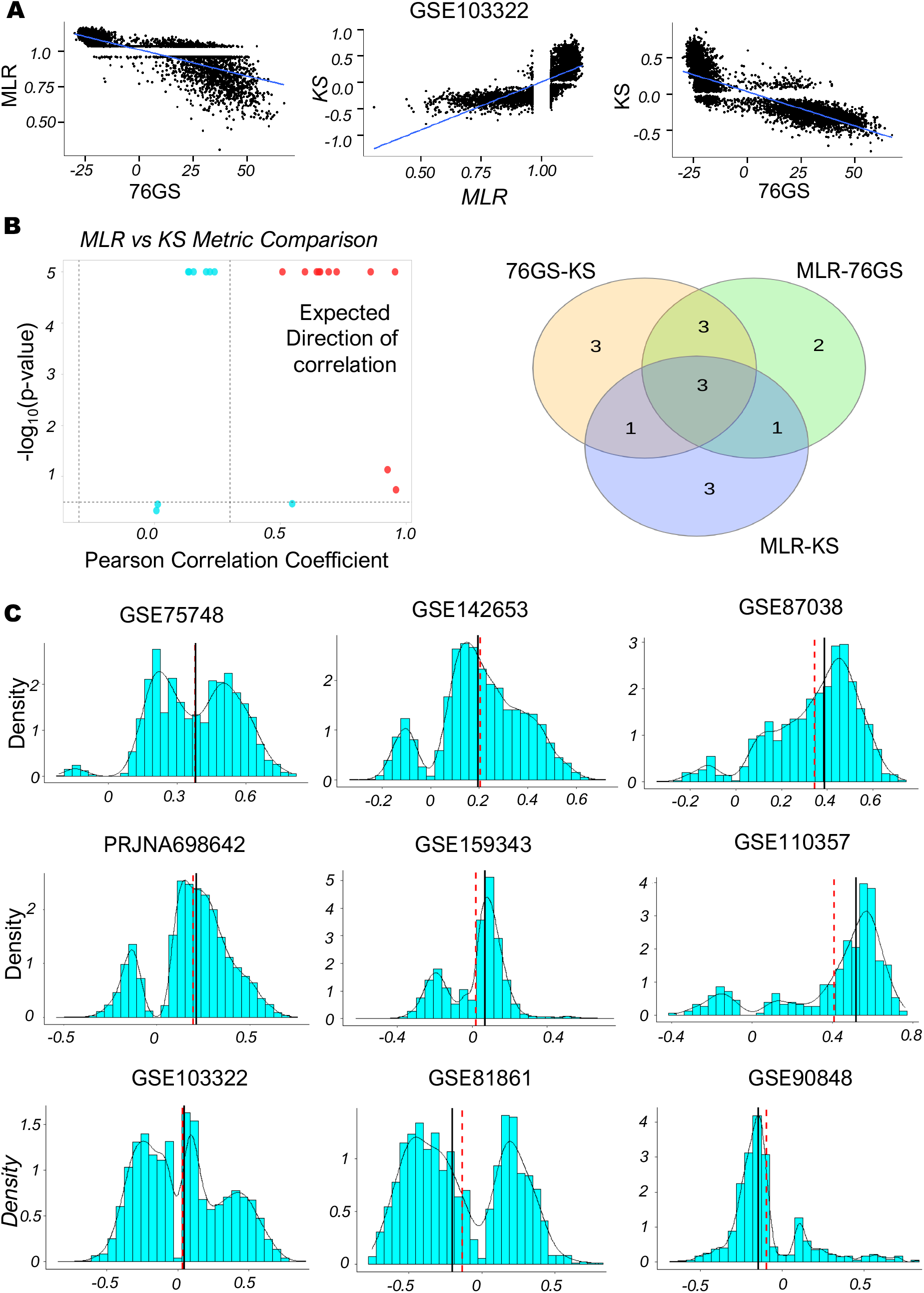
Concordance across EMT scoring methods in single cell RNA-Seq datasets. **A)** Scatter plot of pairwise correlation estimated by linear regression (blue) in GSE103322 dataset. **B**) (Left) Plots showing the correlation of MLR vs. KS EMT scoring metrics for different single-cell RNA-Seq datasets, for each sample –log_10_(p-value) is plotted as a function of Pearson’s correlation coefficient. Thresholds for correlation (R < −0.3 or R > 0.3; vertical dashed grey lines) and p-values (p < 0.05; horizontal dashed grey lines) are indicated. (Right) Venn diagram depicting the common datasets across three pairwise comparisons that are significantly correlated in the expected direction. **C)** Density plots and histograms of KS scores in various datasets. Red dashed line and black solid line depicts mean and median respectively.

Next, we plotted the histograms for EMT scores of various single-cell RNA-seq datasets to decipher the heterogeneity seen along the EMP spectrum across a variety of biological contexts: 1. human embryonic stem-cell-derived progenitor cells differentiating to endoderm (GSE75748) (Chu et al., 2016); 2. human fetal pituitary gland development including progenitors of many endocrine cell types and subtypes (GSE142653) (Zhang et al., 2020); 3. cells from different tissues and organs of E9.5 to E11.5 mouse embryos (GSE87038) (Dong et al., 2018); 4. MCF10A cells treated with TGF-β for varying durations and exhibiting a gradual change in their EMT status (PRJNA698642) (Deshmukh et al., 2021); 5. murine pancreatic duct cells with variations along the EMP spectrum (GSE159343) (Hendley et al., 2021); 6. EpCAM+ and EpCAM-squamous skin carcinoma cells with varied epithelial and/or mesenchymal features (GSE110357) (Pastushenko et al., 2018); 7. cells from oral cavity tumors/head and neck squamous cell carcinoma (GSE103322) (Puram et al., 2017); 8. human colorectal cancer cell lines and tumors (GSE81861) (Li et al., 2017), and; 9. mouse hair follicle stem cells and transit-amplifying cells (GSE90848) (Yang et al., 2017). Across these cases, we observed two distinct peaks in KS scoring metrics (**Fig 3C**), suggesting the presence of at least two major subpopulations with varied EMT status.

Of note, several plots for the 76GS and MLR metrics appeared saturated, which we hypothesized related to the relative sparsity of predictor signal in the single cell datasets. For the MLR approach, we then restricted our analysis to datasets with at least 90% of all single-cell samples containing nonzero entries for each predictor, indicating the presence of measurable signal. In these cases, MLR and 76GS metrics were able to recapitulate the trends observed in KS for many such datasets (**Fig S5A-B**).

### Quantifying the EMP spectrum during lung diseases and cellular reprogramming

All the three EMT metrics (76GS, KS, MLR) have been designed and/or trained for quantifying the EMT status in cancer samples (Byers et al., 2013; George et al., 2017; Tan et al., 2014), but our single-cell RNA-seq data analysis suggests their applicability in various developmental contexts. Thus, we investigated if they can be broadly applied to quantifying EMT status in other biological processes other than cancer. For both microarray and RNA-seq datasets (**Table S4**), we used these metrics to calculate EMT status for lung diseases including chronic obstructive pulmonary disease (COPD) and idiopathic pulmonary fibrosis (IPF) where EMT is reported to be involved in initiating and/or aggravating the disease (Jolly et al., 2018).

As compared to normal lung tissues, the fibrotic lung tissues from IPF patients had higher MLR and KS scores but lower 76GS scores, indicating their enhanced mesenchymal status (**Fig 4A, i**; GSE72073). Fibrotic lung tissues had reduced levels of USP13, a deubiquitylase that stabilizes PTEN, and *in vitro* analysis suggested that USP13 deficiency increased invasive and migratory capacities of fibroblasts, traits usually associated with EMT (Geng et al., 2015). Similarly, relative to healthy volunteers, COPD patients showed increased EMT in their bronchoalveolar lavage (BAL) cells (**Fig 4A, ii**; GSE73395). Consistently, as compared to normal lung tissue, patients with any of the three lung pathological situations – IPF, non-specific interstitial pneumonia (NSIP) and mixed IPF-NSIP – exhibited trends of enhanced EMT (**Fig 4A, iii**; GSE110147) (Cecchini et al., 2018). Further, RNA-seq analysis of lung tissues of patients with acute lung injury (ALI) and IPF had higher MLR and KS scores but reduced 76GS scores (**Fig 4A, iv**; GSE134692), consistent with earlier reports (Cabrera-benítez et al., 2012; Gouda et al., 2018; Sivakumar et al., 2019).

**Figure 4.**
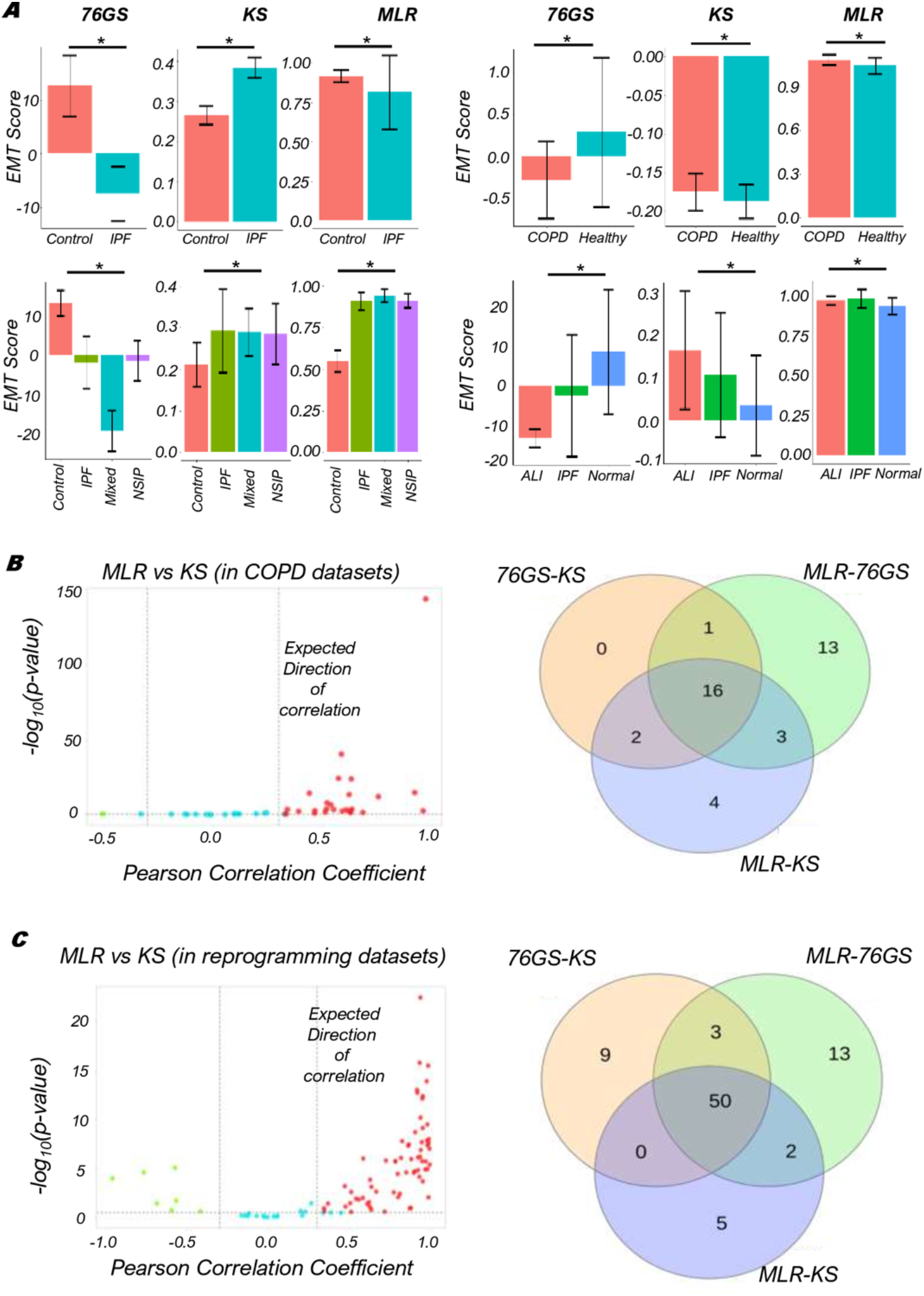
Concordance across all three EMT scoring methods in COPD/IPF and reprogramming datasets. **A)** Bar plots showing EMT scores of different datasets calculated using three EMT scoring methods – i) Lung tissues of healthy cases and IPF patients (GSE72073), ii) EMT status in BAL cells of healthy volunteers and COPD patients (GSE73395), iii) Lung tissues of IPF, NSIP and Mixed IPF/ NSIP patients, as well as healthy lung tissue (GSE110147), iv) EMT status in lung tissues of Normal, ALI and IPF patients (GSE134692). **B)** (Left) Plots of the correlation of MLR vs KS EMT scoring metrics across 46 datasets. For each sample, –log10(p-value) is plotted as a function of Pearson’s correlation coefficient. Thresholds for correlation (R < −0.3 or R > 0.3; vertical dashed grey lines) and p-values (p < 0.05; horizontal dashed grey lines) are shown. (Right) Venn diagram depicting the common datasets across all pairwise comparisons that are significantly correlated in the expected direction. **C)** Same as B) but for reprogramming associated datasets.

After investigating these few examples, we analyzed the trends among KS, MLR and 76GS scores obtained for 46 microarray or RNA-seq datasets associated with lung injury. Reinforcing the trends seen for cancer-related datasets, 76GS and MLR scores were negatively correlated in roughly 72% (33/46) of datasets. Similarly, 76GS and KS scores correlated negatively in ~41% (19/46) of datasets. Further, KS and MLR scores correlated positively in ~54% (25/46) of datasets (**Fig 4B, S4B**). Overall, we see a strong concordance among the three EMT scoring metrics for non-cancerous lung diseases too.

Further, we investigated a set of datasets related to cellular reprogramming of differentiated cell types to induced pluripotent stem cells (iPSCs), where EMT/MET are reportedly involved (Lai et al., 2020) (**Table S5**). Across 92 datasets for which we calculated the 76GS, KS and MLR EMT scores, roughly 62% (57/92) showed a positive correlation between MLR and KS, while 67% (62/92) showed negative correlation between KS and 76GS, and approximately 74% (68/92) showed a negative correlation between corresponding 76GS and MLR ones. Overall, 54% (50/92) datasets demonstrated all three pairwise correlations to be strong (**Fig 4C, S4C**) thus endorsing that these EMT scoring metrics can be quite consistent with one another in terms of identifying the EMP status of cells *en route* to cellular reprogramming.

### Context-specific association of EMP status with patient survival

Next, we quantified EMT scores in patient samples using TCGA datasets of various cancer types. Here also we found the expected trends that the 76GS scores shows negative correlation with the MLR and KS scores and KS and MLR scores are positively correlated to each other (**Fig 5A**), reinforcing our observations for a pan-cancer analysis of microarray datasets (**Table S6**) (Vasaikar et al., 2021). We also calculated Single-set Gene Set Enrichment Analysis (ssGSEA scores) (Subramanian et al., 2005) using the EMT gene set from MSigDB (Liberzon et al., 2011). Each ssGSEA enrichment score represents the degree to which the genes in a particular gene set are coordinately regulated within a sample. We find that the ssGSEA scores for EMT show, as expected, negative correlation with 76GS scores and positive correlation with MLR and KS scores.

**Figure 5.**
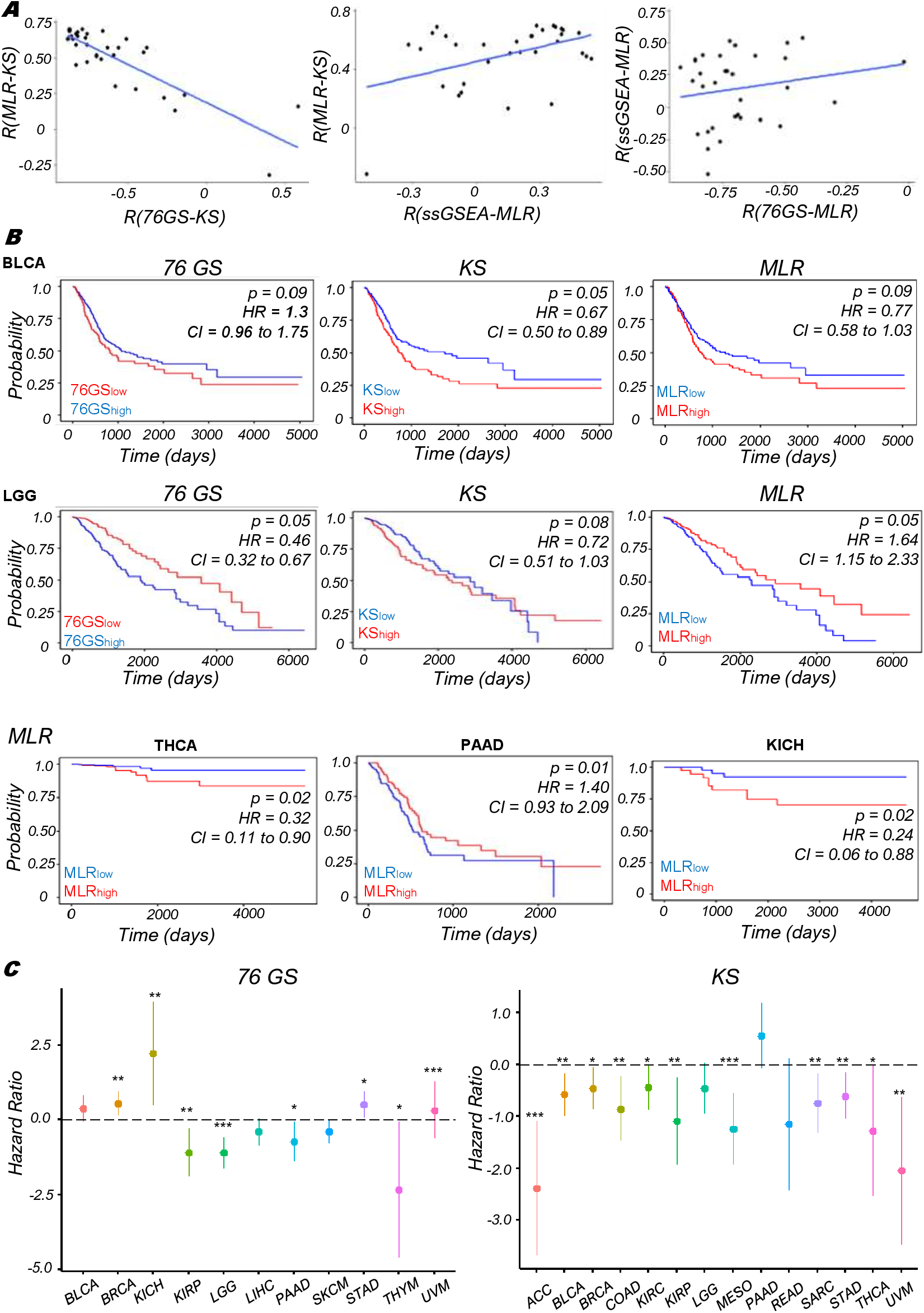
EMT scoring in TCGA datasets and survival analysis using patient samples. **A)** Scatter plot of pairwise correlation estimated by linear regression (blue) in TCGA datasets. **B)** Correlation between EMT score (high vs. low) and overall survival (OS) in various TCGA datasets. Kaplan–Meier survival analysis is performed to estimate differences in survival for different EMT metrics. p-values (p) reported are based on log rank test. The values of Hazard ratio (HR) with 95% confidence interval (CI) values are included. **C)** Plot of log_2_ hazard ratio (HR; mean ± 95% confidence interval) comparing overall survival (OS) of 76GS (left) and KS (right) EMT scoring metrics on different TCGA cancers and cohorts. p-values are based on log-rank test, and those with significant differences (p < 0.05) are marked with a star (*).

We also assessed the association between EMT scores and patient survival using different survival data types (Overall survival (OS), Disease-Specific Survival (DSS), Progression free interval (PFI) and Disease-free interval (DFI)) in various TCGA cancer cohorts. The samples were scored using all three methods and segregated into high and low groups based on the mean value of each EMT score. The 76GS^low^ subgroup can be thought of as comparable to the MLR^high^ and KS^high^ groups, given their relatively strong M signature. In bladder cancer (BLCA; **Fig 5B**, top), we see consistent trends in case of overall survival (OS) for all three types of EMT scores that the stronger the M phenotype, poorer the survival probability; whereas in case of Low-Grade Glioma (LGG), we see the opposite trend, that is, the stronger the M phenotype, the better the survival probability for 76GS and MLR (**Fig 5B**, middle). Similarly, in Thyroid Cancer (THCA) and Kidney Chromophobe (KICH), higher MLR scores reflect worse survival outcomes, but in pancreatic adenocarcinoma (PAAD), higher MLR scores associate with better outcomes (**Fig 5B**, bottom), indicating a context-specific association of the extent of EMT with patient survival. These trends for OS were also seen in 76GS and KS scores, where the hazard ratio (HR) > 1 and HR < 1 scenarios were both observed (**Fig 5C, S6**) depending on the cancer subtype in TCGA.

After investigating OS data, we calculated the survival probabilities through other metrics as well – DSS, PFI and DFI, wherever available. For DSS, we found that Kidney Papillary Cell Carcinoma (KIRP) samples having larger MLR scores (or lower 76GS scores) corresponding reflect poorer survival. This trend held for PFI as well with high KS scores reflecting poorer survival (**Fig 6A**; columns 1, 2), but contrasted with LGG samples, which indicated the same trend of improved DSS and PFI associated with higher (resp. lower) MLR (resp. 76GS) scores (**Fig 6B**; columns 1, 2). The DFI for Head and Neck Cancer (HNSC) indicates a worse prognosis associated with high MLR scores, opposite to that seen for the case of Uterine Carcinosarcoma (UCS) (**Fig 6C**), indicating that the UCS samples with enriched M phenotypes correspond to improved survival.

**Figure 6.**
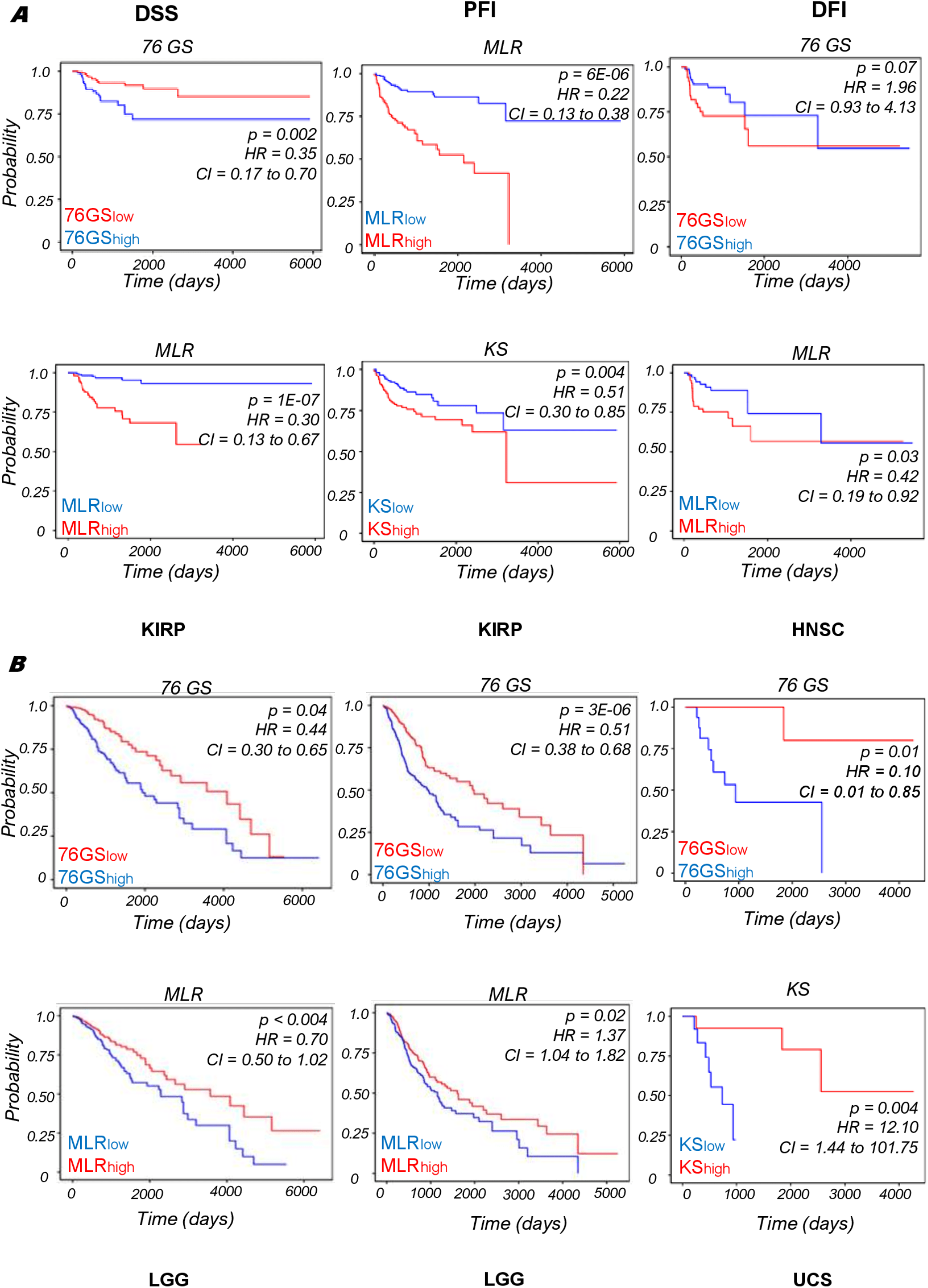
Correlation between EMT score (high vs. low) and various survival types (DSS, PFI and DFI) in various TCGA datasets. Kaplan–Meier survival analysis is performed to estimate differences in survival for different EMT metrics. p-values (p) reported are based on the log rank test. The values of Hazard ratio (HR) with 95% confidence interval (CI) values are included

## Discussion

Quantifying the spectrum of epithelial-hybrid-mesenchymal cell states in cancer has garnered recent interest due to a surge in the availability of *in vitro* and *in vivo* spatial and/or temporal dynamic and high-throughput data at multiple levels - transcriptomic, proteomic, epigenetic, metabolic, and morphological (Bocci et al., 2019; Cook and Vanderhyden, 2020; Deshmukh et al., 2021; Devaraj and Bose, 2019; Jia et al., 2019; Johnson et al., 2021; Karacosta et al., 2019; McFaline-Figueroa et al., 2019; Serresi et al., 2021; Stylianou et al., 2019; Wang et al., 2020). Phenotypic plasticity and heterogeneity along the EMP spectrum has been postulated to be a more important criteria for defining the survival fitness of a cancer cell population than the predominance of a specific phenotype (Brown et al., 2021; Chakraborty et al., 2021), suggesting possible benefits to a more heterogeneous population through cooperation among cancer cells with varying EMP phenotypes (Neelakantan et al., 2017; Tsuji et al., 2008). For single cells, hybrid E/M phenotypes are believed to be the most plastic relative to their more ‘extreme’ epithelial and mesenchymal counterparts; such plasticity can amplify tumor-initiation ability (Kröger et al., 2019; Ruscetti et al., 2016). Therefore, characterizing the EMP as a continuous spectrum instead of as an ‘all-or-none’ process becomes imperative for an improved understanding of emergent dynamics of EMT and MET, and their relevance to patient survival.

Here, we used three different EMT transcriptomic-based scoring metrics, each of which was developed using cancer cell lines and/or tumor samples, to quantify the extent of EMT on a continuum – 76GS, KS, MLR. Using these metrics, we calculated EMT scores for over 100 RNA-seq datasets – both at bulk and individual cell levels – across multiple biological contexts (cancer, fibrosis, COPD, and cellular reprogramming to iPSC). We observed that these methods show a high degree of concordance among themselves in their ability to identify the extent of EMT/MET a sample has undergone, despite using different gene lists and algorithms. This concordance suggests an overlap of core expression patterns central to EMT in a high-dimensional feature space and indicates that these metrics – initially developed for cancer samples – can be applied more generally to a broader range of biological contexts. Using these metrics in biological contexts where hybrid E/M states have been proposed (Aban et al., 2020; Grande et al., 2015; Jolly et al., 2018) may be helpful in mapping the corresponding trajectories of EMT/MET. The hybrid E/M state has been previously documented at both bulk and single-cell levels during various stages of development as well (Dong et al., 2018; Leroy and Mostov, 2007); here, we have shown proof-of-principle that these scoring metrics can successfully quantify the extent of EMP in mouse models. However, whether they can be adapted to adequately investigate the role of EMT in applications for other non-human model organisms remains to be investigated.

In our analysis of single-cell RNA-seq data, the resolvability of bimodal distributions consistent with dual sub-populations was optimally characterized by the KS scoring metric. Improvement in the MLR and 76GS approaches were observed when restricting the analysis to datasets having non-zero MLR predictors for a majority (>90%) of single-cell samples. These scores, while concentrated in the middle of the EMT interval, were able to recover features of the distributions observed via the KS method. Together, this suggests that the development of optimized signal-to-noise criteria, may improve the absolute placement of samples on the EMT MLR spectrum and is the focus of research effort. Future efforts should consider how these metrics can be adapted to investigate different cell-state transition trajectories, for instance, by defining a two-dimensional EMT score that can deconvolute gains in mesenchymal program vs. losses in epithelial one (Foroutan et al., 2018).

These metrics have been helpful in investigating the association of EMT/MET with other axes of cellular plasticity such as stemness (Bocci et al., 2018), immune evasion (Li et al., 2019) and sensitivity to anti-cancer agents (Wang et al., 2021). Intriguingly, EMT status of primary tumors was not found to be universally correlated with worse patient survival, but instead showed a context-dependent trend, consistent with previous reports (Tan et al., 2014). EMP is a highly dynamic trait. Thus, capturing static snapshots of gene expression profiles may not be sufficient for recapitulating the dynamic dependence of cancer cell fitness on EMT and/or MET. Thus, the EMT status and/or heterogeneity of a primary tumor may not reflect that of circulating tumor cells (CTCs) and their metastatic potential, leading to such observed context-specific trends. Moreover, transcriptomic profiles may not be sufficient to indicate phenotypic variability and incorporate epigenetic and/or metabolic status can elucidate the manifestations of dynamic adaptation during metastasis. Understanding the interplay among EMT, metabolic and epigenetic reprogramming (Dumont et al., 2008; Jia et al., 2021, 2019; Peixoto et al., 2019; Serresi et al., 2021) will be key for better patient stratification and therapeutic strategies.

## Materials and Methods

### Software and Datasets

We downloaded high-throughput transcriptomics data (bulk and single cell) from GEO and EMBL-EBI databases. Microarray datasets were downloaded using GEOquery R Bioconductor package (Davis and Meltzer, 2007). TCGA expression and survival data were obtained from the UCSC xena browser (https://xena.ucsc.edu/). Statistical analysis, survival analysis and plots were all done in R version 4.0.3. ggplot was used for plots.

### Preprocessing of datasets

After downloading the HTS datasets, quality check was done by FASTQC (Andrews, 2010) (https://www.bioinformatics.babraham.ac.uk/projects/fastqc/). Bulk and single-cell RNA seq data were aligned to reference genome (hg38/mm10, appropriately) using STAR-aligner (Dobin et al., 2013). Samtools (Li et al., 2009) was used to modify alignment files (SAM/BAM) and htseq-count (Anders et al., 2015) was used to calculated the read counts. Using these read counts, TPM expression was calculated using custom scripts and log2 normalized TPM values were used for calculation of EMT scores. In case of microarray datasets, they were preprocessed to obtain the gene-wise expression for each sample from probe-wise expression matrix. If there were multiple probes mapping to one gene, then the mean expression of all the mapped probes was considered for that gene.

### Calculation of EMT scores

EMT scores were calculated using all three methods - 76GS, KS and MLR as previously done for microarray datasets (Chakraborty et al., 2020). MLR method, which was designed for microarray datasets (George et al., 2017), was adjusted to work for HTS transcriptomics data as well.

### MLR model applied to RNA-Seq

We adapted a previously developed method of quantifying EMT spectrum trained on and designed to predict microarray samples (George et al., 2017). Very briefly, VIM, CDH1, and CLDN7 transcripts were identified to maximally predict NCI-60 holdout samples in leave-one-out assessment utilizing two-dimensional multinomial logistic regression (MLR). These, together with a list of 20 normalizers, enable the assignment of each input (CLDN7, VIM/CDH1) to an ordered triple (P_E_, P_E/M_, P_M_) that characterizes the probability that a signature belongs either to the Epithelial (E), Mesenchymal (M), or hybrid (E/M) group. This ordered triplet is then projected onto the interval [0, 2] with 0 designating a fully epithelial signature, 1 maximally hybrid signature, and 2 fully mesenchymal signature.

In order to apply the microarray based MLR model to RNA-seq data, we utilized transcriptomic data available in both formats on biological replicates (Zhao et al., 2014). Using the data available in Fig. 2 (2 biological replicates for each of 6 time points), we restricted our analysis to the intersection of microarray and RNA-seq transcripts for genes represented in positive abundance for both datasets. Linear regression to the average of each biological replicate, producing a total of 6 slope-intercept pairs:

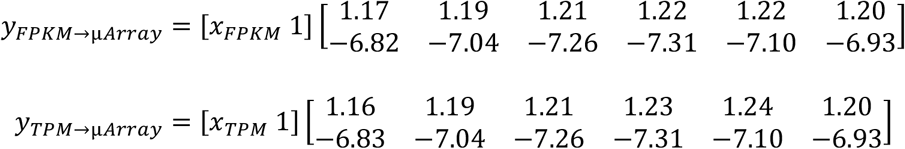

which were then averaged to be used as the fit parameters for cross-platform assessment:

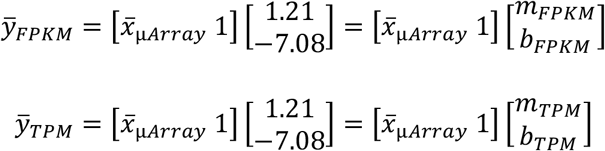

From these, unique microarray values, *x*_μ*Array*_, representative of RNASeq values may be calculated by inversion:

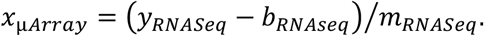

### Survival Analysis

Different metrics of survival data was obtained from TCGA cohort. All samples were divided into 76GS^high^ and 76GS^low^, MLR^high^ and MLR^low^, KS^high^ and KS^low^ groups based on the mean (or median) of the respective scores of the samples. Kaplan–Meier analysis was done using R package “survival” and plotted using R package “ggfortify”. Log rank test was used to calculate the p-values. Hazard ratio (HR) and confidence interval (95% CI) reported are estimated using cox regression.

### T-test

Two-tailed student’s t-test with unequal variance was done to compare between samples in many bar plots. Error bars denoted the standard deviation (statistical significance at p < 0.05).

### ssGSEA

Single-sample GSEA (ssGSEA), an extension of Gene Set Enrichment Analysis (GSEA), calculates separate enrichment scores for each pairing of a sample and gene set. Each ssGSEA enrichment score represents the degree to which the genes in a particular gene set are coordinately up- or down-regulated within a sample. We used “HALLMARK_EMT” gene set from The Molecular Signatures Database (MSigDB) database and the scores were calculated using R package “ssgsea”.

## Code availability

All codes used in the manuscript are given at https://github.com/sushimndl/EMT_Scoring_RNASeq

## Acknowledgements

MKJ was supported by Ramanujan Fellowship awarded by SERB, DST, Government of India (SB/S2/RJN-049/2018) and by the InfoSys Young Investigator Fellowship awarded by InfoSys Foundation, Bangalore. SSS is supported by grants from Clifford Craig Foundation Launceston General Hospital and Rebecca L Cooper Medical Research Foundation.

## Author contributions

MKJ, JTG, and HL conceived of and designed the research; MKJ and JTG supervised the research; SM, TT, RJ, SS and PT performed the research. SM, TT, SS, PC and SSS analyzed and interpreted the data. All authors contributed to manuscript writing.

## Supplementary Figures

**Figure S1.**
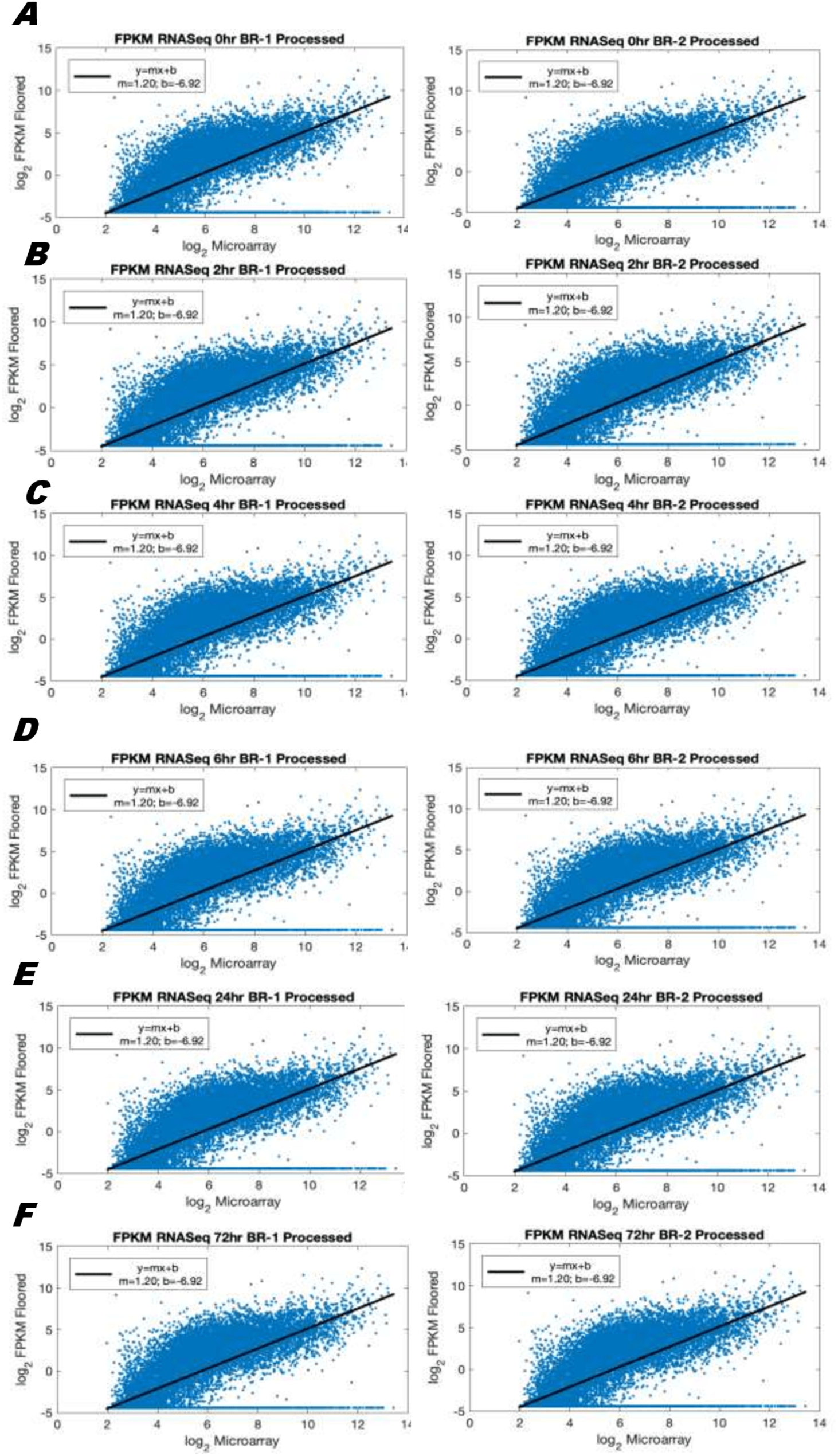
Linear regression on log_2_ FPKM-normalized RNA-seq and log_2_ microarray correlation plots. Data from Fig. 2 of (Zhao et al., 2014) was plotted for each of 2 biological replicates for A. 0hr; B. 2hr; C. 4hr; D. 6hr; E. 24hr; F. 72hr timepoints. Slope-intercept parameters are reported in the figure legend of each regression plot.

**Figure S2.**
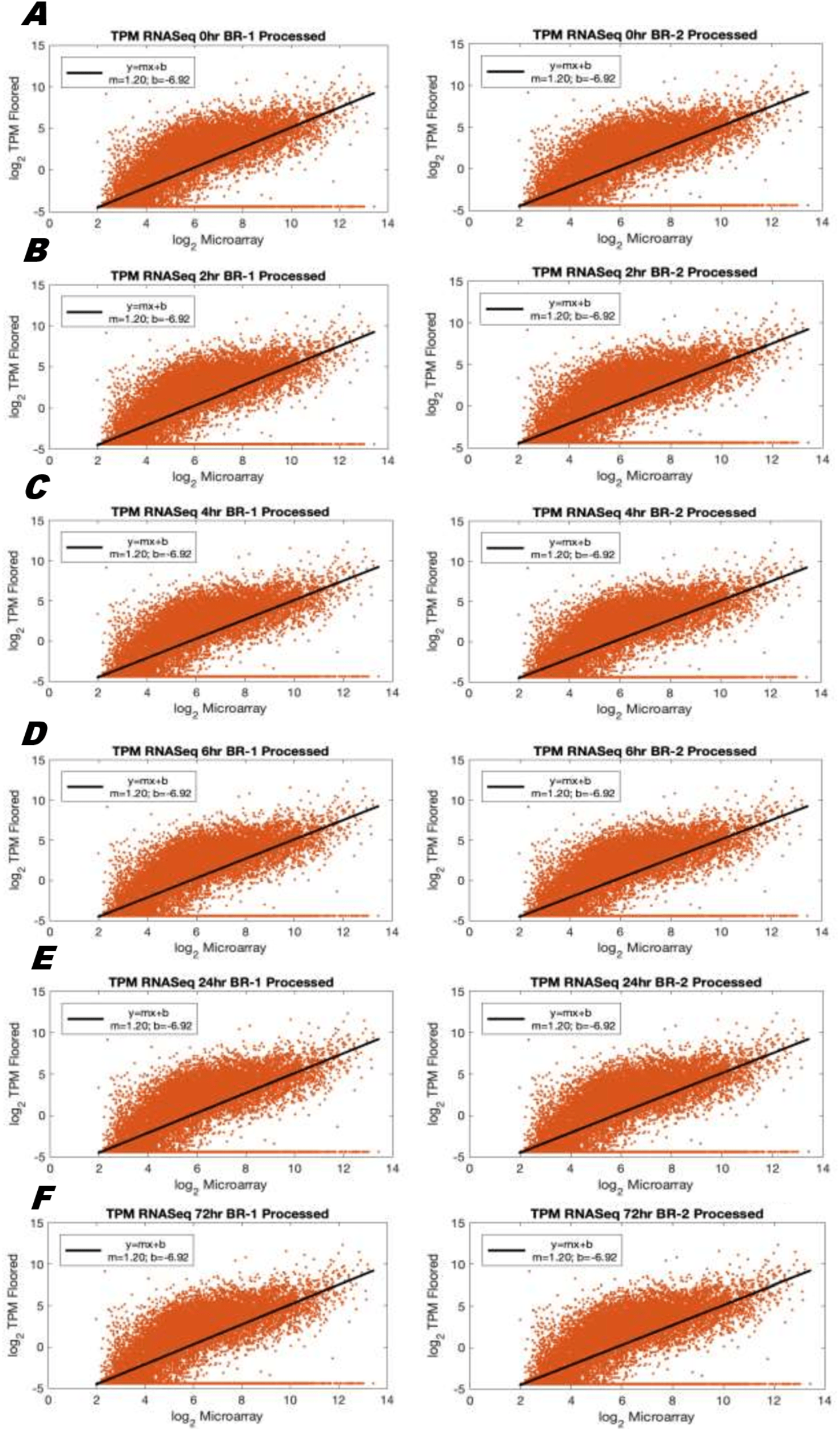
Linear regression on log_2_ TPM-normalized RNA-seq and log_2_ microarray correlation plots. Data from Fig. 2 of (Zhao et al., 2014) was plotted for each of 2 biological replicates for A. 0hr; B. 2hr; C. 4hr; D. 6hr; E. 24hr; F. 72hr timepoints. Slope-intercept parameters are reported in the figure legend of each regression plot.

**Figure S3.**
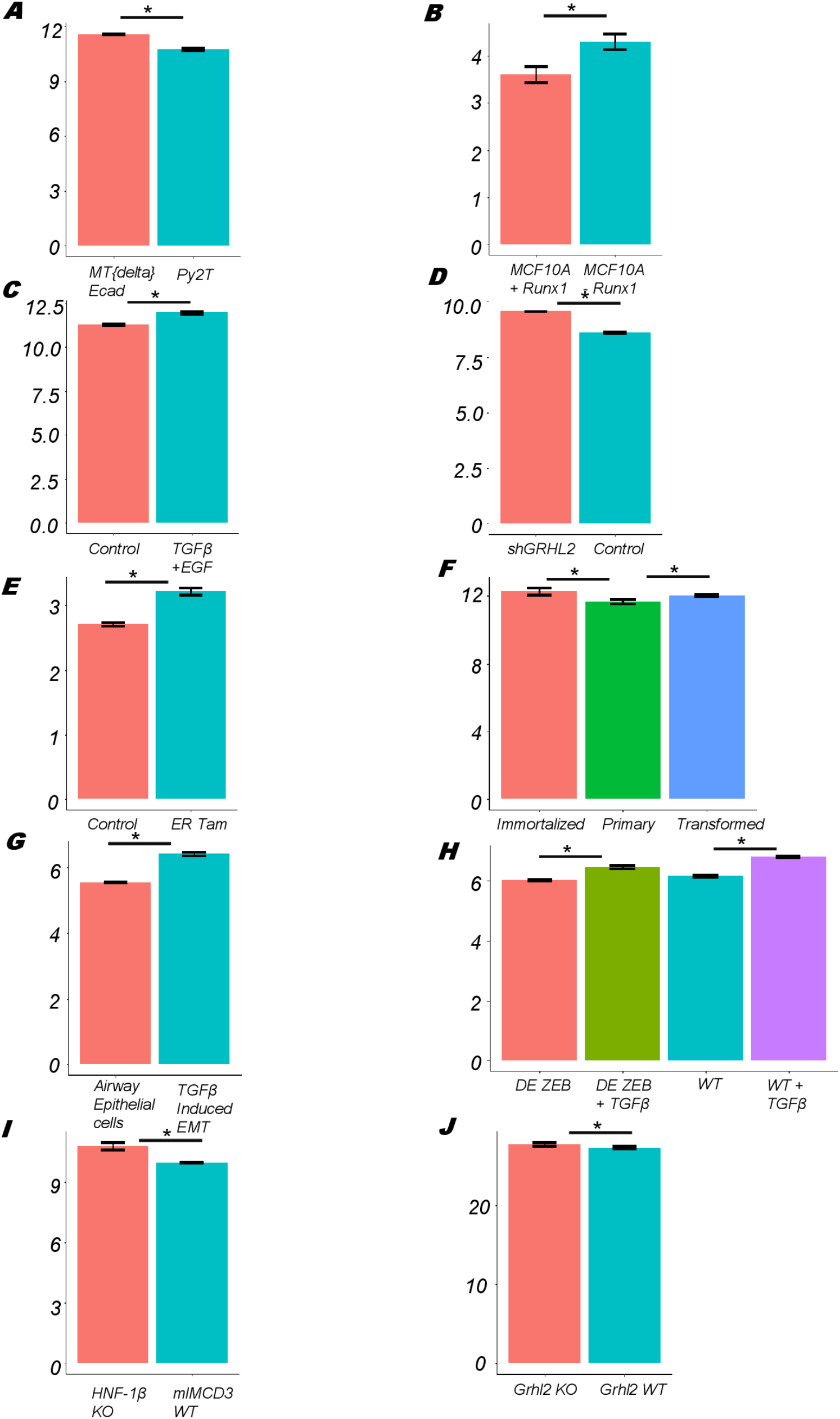
Bar plots showing ssGSEA scores of different datasets (ref Fig 2) calculated using EMT gene set from MSigDB. **A)** GSE118612 **B)** GSE85857 **C)** GSE72419 **D)** GSE118407 **E)** GSE139074 **F)** GSE110677 **G)** GSE61220 **H)** GSE124843 **I)** GSE97770 **J)** GSE106130

**Figure S4.**
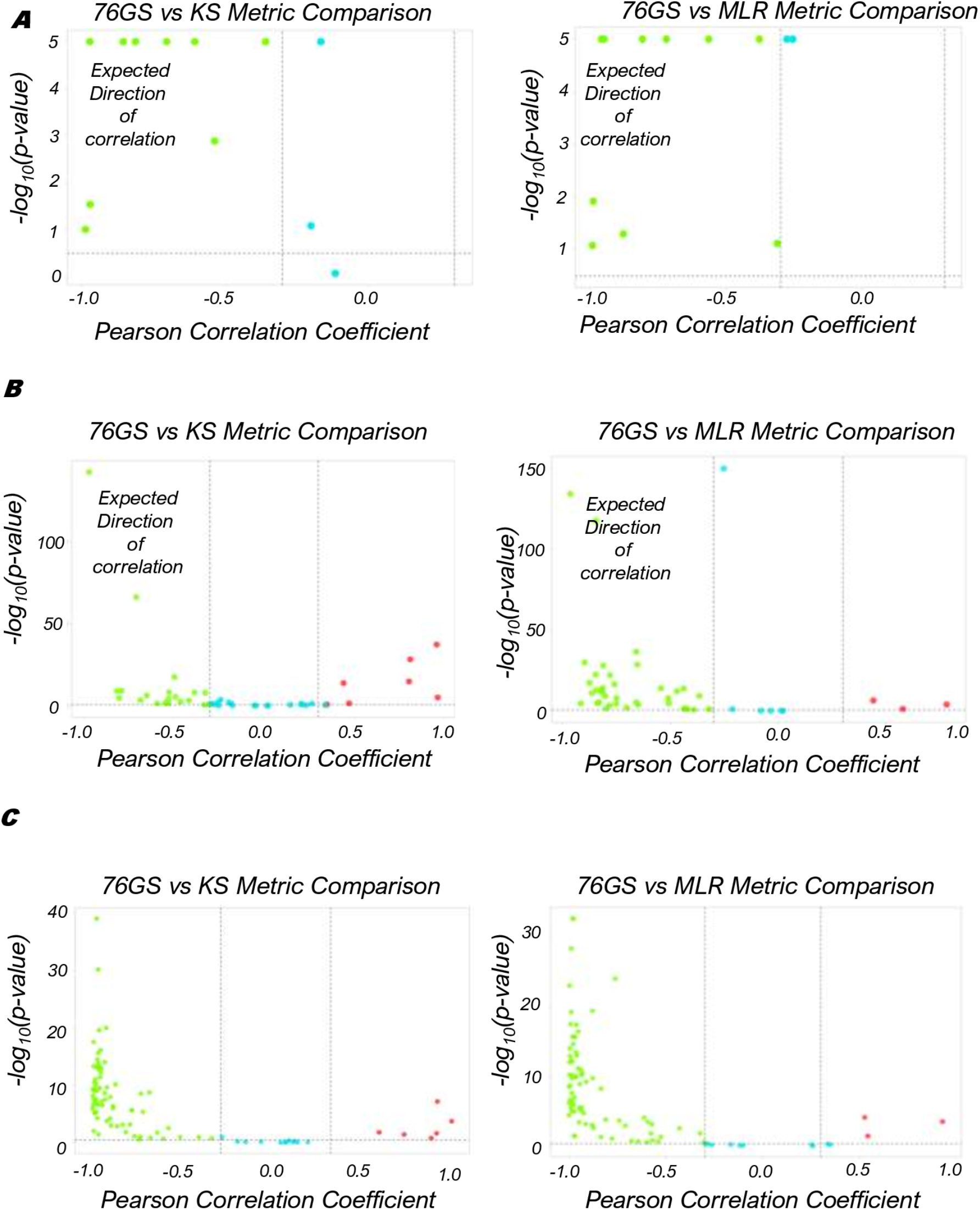
Concordance among KS, 76GS and MLR metrics in multiple biological contexts. **A, B)** Plots showing the correlation of EMT scoring metrics across single-cell RNA-seq data, −log_10_ (p-value) is plotted as a function of Pearson’s correlation coefficient. Thresholds for correlation (R < −0.3 or R > 0.3; vertical dashed grey lines) and p-values (p < 0.05; horizontal dashed grey lines) are denoted. **C, D)** Same as A, B but for datasets pertaining to lung diseases (COPD, IPF etc.). **E, F)** Same as A, B but for datasets pertaining to cellular reprogramming to iPSCs.

**Figure S5.**
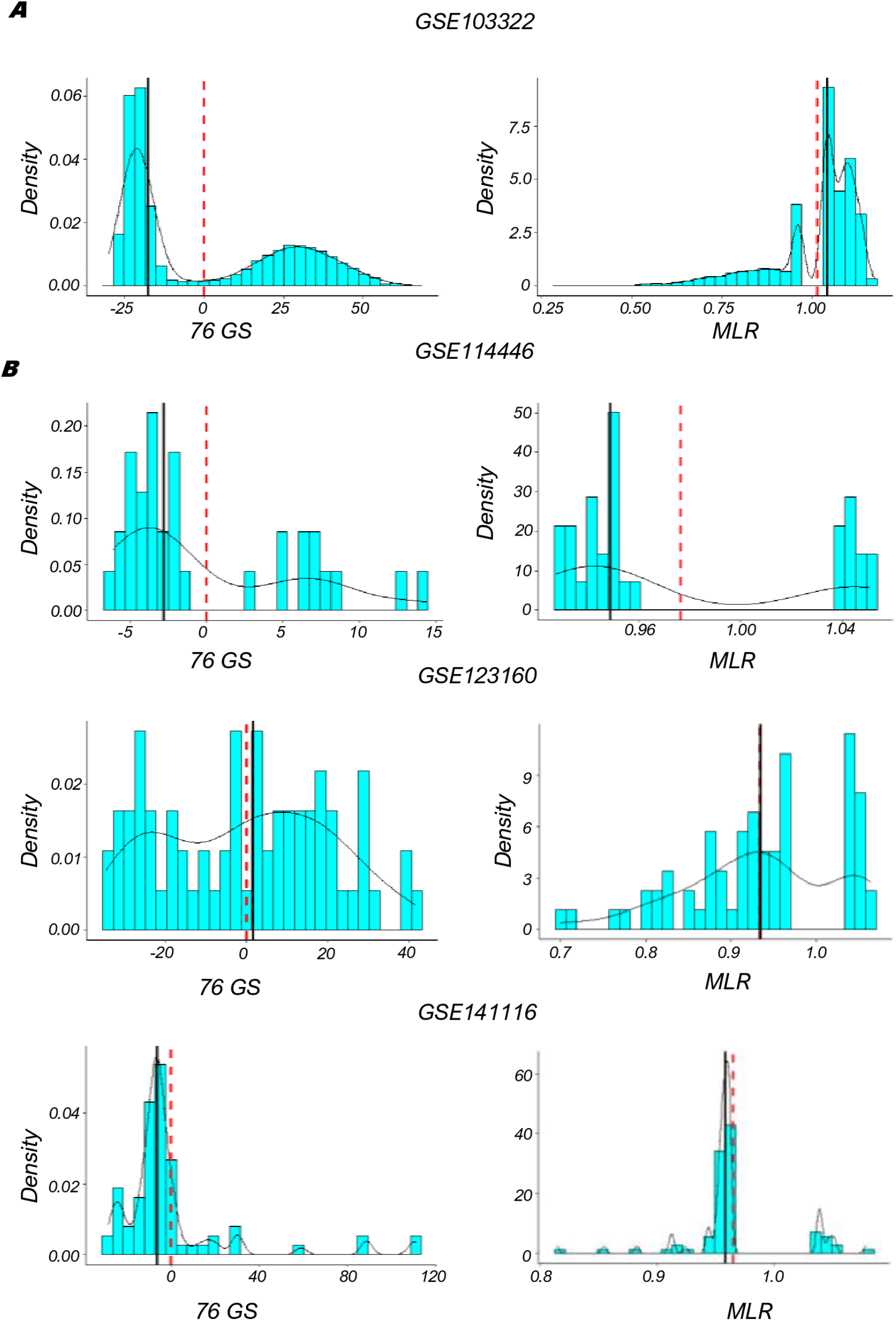
Heterogeneity in EMP. Density plots and histograms of MLR and KS scores in various datasets as indicated. Red dashed line and black solid line depicts mean and median respectively.

**Figure S6.**
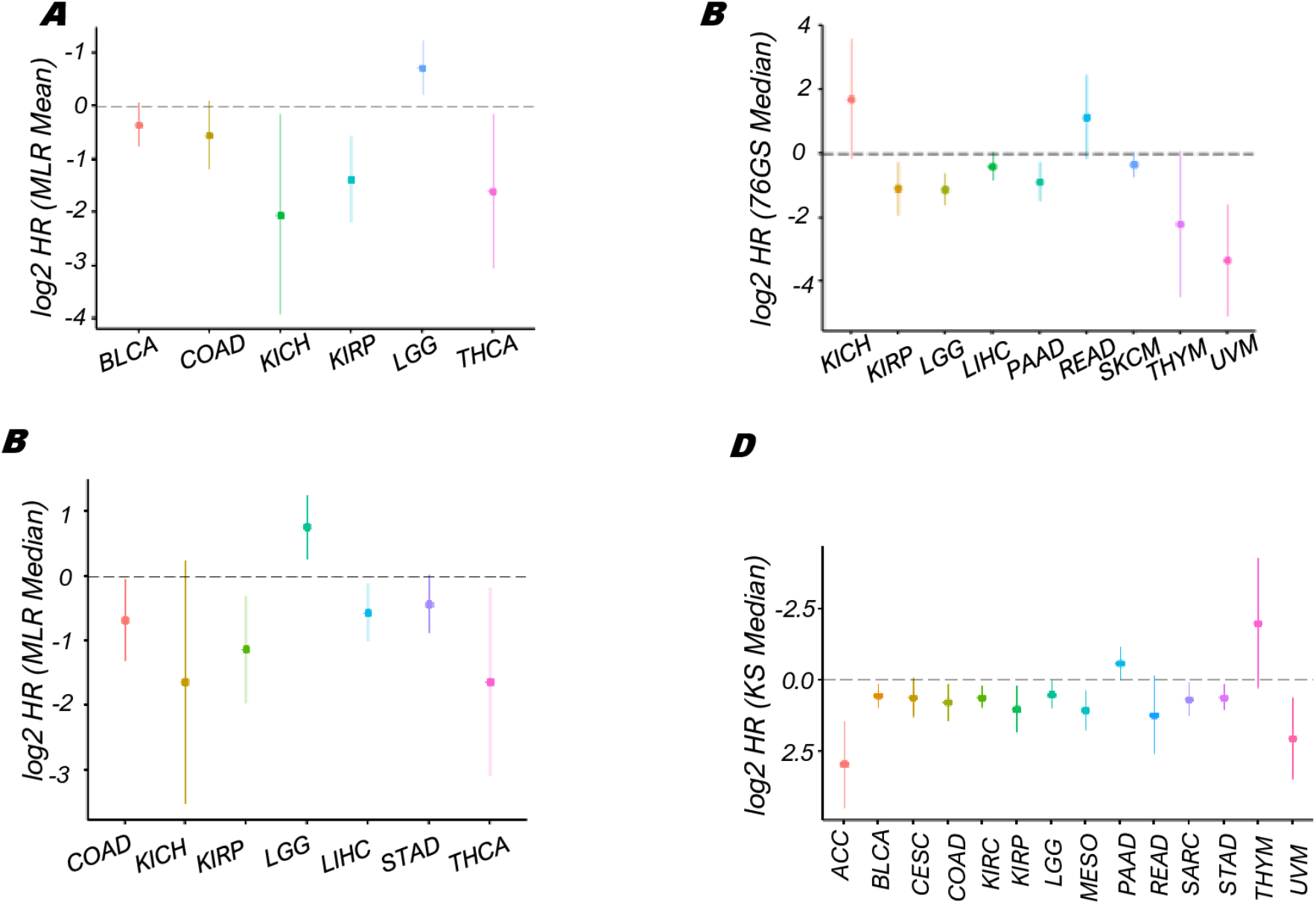
Kaplan-Meier analysis. Plot of log2 hazard ratio (HR) with 95% confidence interval, comparing the overall survival (OS) of 76GS (left) and KS (right) EMT scoring metrics on different TCGA cancer types, where samples are segregated either by mean or by median.

## Legends for Supplementary Tables

**Table S1**. Details of 77 bulk RNA-seq datasets (GSE ID, PMID, no. of samples, and pairwise correlation coefficient and p-values).

**Table S2**. Details of sample IDs in individual datasets that have been considered for comparative analysis in Fig 2.

**Table S3.** Details of 19 single-cell RNA-seq datasets (GSE ID, PMID, no. of samples, and pairwise correlation coefficient and p-values).

**Table S4.** Details of 46 RNA-seq and microarray datasets pertaining to COPD, IPF and other lung diseases (GSE ID, PMID, no. of samples, and pairwise correlation coefficient and p-values).

**Table S5.** Details of 92 RNA-seq and microarray datasets pertaining to cellular reprogramming (GSE ID, PMID, no. of samples, and pairwise correlation coefficient and p-values).

**Table S6.** Details of datasets included in EMT-ome (Vasaikar et al., 2021) together with pairwise correlation coefficient and p-values among EMT scoring metrics.

## References

Aban, C., Lombardi, A., Neiman, G., Biani, M.C., La Greca, A., Walsman, A., Moro, L.N., Sevlever, G., Miriuka, S., Luzzani, C., 2020. Downregulation of E-cadherin in pluripotent stem cells triggers partial EMT. bioRxiv 101899. doi:10.1101/2020.05.18.101899

Aceto, N., Bardia, A., Miyamoto, D.T., Donaldson, M.C., Wittner, B.S., Spencer, J.A., Yu, M., Pely, A., Engstrom, A., Zhu, H., others, 2014. Circulating tumor cell clusters are oligoclonal precursors of breast cancer metastasis. Cell 158, 1110–1122.

Anders, S., Pyl, P.T., Huber, W., 2015. HTSeq-A Python framework to work with high-throughput sequencing data. Bioinformatics 31, 166–169. doi:10.1093/bioinformatics/btu638

Andrews, S., 2010. FastQC: a quality control tool for high throughput sequence data.

Aue, A., Hinze, C., Walentin, K., Ruffert, J., Yurtdas, Y., Werth, M., Chen, W., Rabien, A., Kilic, E., Schulzke, J.D., Schumann, M., Schmidt-Ott, K.M., 2015. A grainyhead-like 2/Ovo-like 2 pathway regulates renal epithelial barrier function and lumen expansion. J. Am. Soc. Nephrol. doi:10.1681/ASN.2014080759

Bierie, B., Pierce, S.E., Kroeger, C., Stover, D.G., Pattabiraman, D.R., Thiru, P., Liu Donaher, J., Reinhardt, F., Chaffer, C.L., Keckesova, Z., Weinberg, R.A., 2017. Integrin-β4 identifies cancer stem cell-enriched populations of partially mesenchymal carcinoma cells. Proc. Natl. Acad. Sci. 114, E2337–2346. doi:10.1073/pnas.1618298114

Bocci, F., Gearhart-Serna, L., Boareto, M., Riberio, M., Ben-Jacob, E., Devi, G.R., Levine, H., Onuchic, J.N., Jolly, M.K., 2019. Toward understanding cancer stem cell heterogeneity in the tumor microenvironment. Proc Natl Acad Sci U S A 116, 148–157. doi:10.1073/pnas.1815345116

Bocci, F., Jolly, M.K., George, J.T., Levine, H., Onuchic, J.N., 2018. A mechanism-based computational model to capture the interconnections among epithelial-mesenchymal transition, cancer stem cells and Notch-Jagged signaling. Oncotarget 9, 29906–20. doi:10.18632/oncotarget.25692

Bocci, F., Mandal, S., Tejaswi, T., Jolly, M.K., 2021. Investigating epithelial-mesenchymal heterogeneity of tumors and circulating tumor cells with transcriptomic analysis and biophysical modeling. Comput. Syst. Oncol. in press. doi:10.1002/cso2.1015

Brabletz, S., Brabletz, T., 2010. The ZEB/miR-200 feedback loop--a motor of cellular plasticity in development and cancer? EMBO Rep. 11, 670–7. doi:10.1038/embor.2010.117

Brown, M.S., Abdollahi, B., Wilkins, O.M., Chakraborty, P., Ognjenovic, N.B., Muller, K.E., Kumar Jolly, M., Hassanpour, S., Pattabiraman, D.R., 2021. Dynamic plasticity within the EMT spectrum, rather than static mesenchymal traits, drives tumor heterogeneity and metastatic progression of breast cancers. bioRxiv 434993. doi:10.1101/2021.03.17.434993

Byers, L.A., Diao, L., Wang, J., Saintigny, P., Girard, L., Peyton, M., Shen, L., Fan, Y., Giri, U., Tumula, P.K., Nilsson, M.B., Gudikote, J., Tran, H., Cardnell, R.J.G., Bearss, D.J., Warner, S.L., Foulks, J.M., Kanner, S.B., Gandhi, V., Krett, N., Rosen, S.T., Kim, E.S., Herbst, R.S., Blumenschein, G.R., Lee, J.J., Lippman, S.M., Ang, K.K., Mills, G.B., Hong, W.K., Weinstein, J.N., Wistuba, I.I., Coombes, K.R., Minna, J.D., Heymach, J. V., 2013. An epithelial-mesenchymal transition gene signature predicts resistance to EGFR and PI3K inhibitors and identifies Axl as a therapeutic target for overcoming EGFR inhibitor resistance. Clin. Cancer Res. 19, 279–290. doi:10.1158/1078-0432.CCR-12-1558

Cabrera-benítez, N.E., Parotto, M., Post, M., Han, B., Spieth, P.M., Cheng, W.-E., Valladares, F., Villar, J., Liu, M., Sato, M., Zhang, H., Slutsky, A.S., 2012. Mechanical stress induces lung fibrosis by epithelial-mesenchymal transition (EMT). Crit. Care Med. 40, 510–517. doi:10.1097/CCM.0b013e31822f09d7.Mechanical

Carpinelli, M.R., de Vries, M.E., Auden, A., Butt, T., Deng, Z., Partridge, D.D., Miles, L.B., Georgy, S.R., Haigh, J.J., Darido, C., Brabletz, S., Brabletz, T., Stemmler, M.P., Dworkin, S., Jane, S.M., 2020. Inactivation of Zeb1 in GRHL2-deficient mouse embryos rescues mid-gestation viability and secondary palate closure. Dis. Model. Mech. 13, dmm042218. doi:10.1242/dmm.042218

Cecchini, M.J., Hosein, K., Howlett, C.J., Joseph, M., Mura, M., 2018. Comprehensive gene expression profiling identifies distinct and overlapping transcriptional profiles in nonspecific interstitial pneumonia and idiopathic pulmonary fibrosis. Respir. Res. 19, 153. doi:10.1186/s12931-018-0857-1

Chakraborty, P., George, J.T., Tripathi, S., Levine, H., Jolly, M.K., 2020. Comparative Study of Transcriptomics-Based Scoring Metrics for the Epithelial-Hybrid-Mesenchymal Spectrum. Front. Bioeng. Biotechnol. 8, 220. doi:10.3389/fbioe.2020.00220

Chakraborty, P., George, J.T., Woodward, W.A., Levine, H., Jolly, M.K., 2021. Gene expression profiles of inflammatory breast cancer reveal high heterogeneity across the epithelial-hybrid-mesenchymal spectrum. Transl. Oncol. 14. doi:10.1016/j.tranon.2021.101026

Chan, S.C., Zhang, Y., Shao, A., Avdulov, S., Herrera, J., Aboudehen, K., Pontoglio, M., Igarashi, P., 2018. Mechanism of Fibrosis in HNF1B-Related Autosomal Dominant Tubulointerstitial Kidney Disease. J Am Soc Nephrol 29, 2493–2509. doi:10.1681/ASN.2018040437

Chen, Limo, Gibbons, D.L., Goswami, S., Cortez, M.A., Ahn, Y.H., Byers, L.A., Zhang, X., Yi, X., Dwyer, D., Lin, W., Diao, L., Wang, J., Roybal, J.D., Patel, M., Ungewiss, C., Peng, David, Antonia, S., Mediavilla-Varela, M., Robertson, G., Jones, S., Suraokar, M., Welsh, J.W., Erez, B., Wistuba, I.I., Chen, Lieping, Peng, Di, Wang, S., Ullrich, S.E., Heymach, J. V., Kurie, J.M., Qin, F.X.F., 2014. Metastasis is regulated via microRNA-200/ZEB1 axis control of tumour cell PD-L1 expression and intratumoral immunosuppression. Nat. Commun. 5. doi:10.1038/ncomms6241

Cheung, K.J., Padmanaban, V., Silvestri, V., Schipper, K., Cohen, J.D., Fairchild, A.N., Gorin, M.A., Verdone, J.E., Pienta, K.J., Bader, J.S., Ewald, A.J., 2016. Polyclonal breast cancer metastases arise from collective dissemination of keratin 14-expressing tumor cell clusters. Proc. Natl. Acad. Sci. 113, E854–E863. doi:10.1073/pnas.1508541113

Chu, L.F., Leng, N., Zhang, J., Hou, Z., Mamott, D., Vereide, D.T., Choi, J., Kendziorski, C., Stewart, R., Thomson, J.A., 2016. Single-cell RNA-seq reveals novel regulators of human embryonic stem cell differentiation to definitive endoderm. Genome Biol. 17, 173. doi:10.1186/s13059-016-1033-x

Chung, V.Y., Tan, T.Z., Tan, M., Wong, M.K., Kuay, K.T., Yang, Z., Ye, J., Muller, J., Koh, C.M., Guccione, E., Thiery, J.P., Huang, R.Y.-J., 2016. GRHL2-miR-200-ZEB1 maintains the epithelial status of ovarian cancer through transcriptional regulation and histone modification. Sci. Rep. 6, 19943. doi:10.1038/srep19943

Chung, V.Y., Tan, T.Z., Ye, J., Huang, R.-L., Lai, H.-C., Kappei, D., Wollmann, H., Guccione, E., Huang, R.Y.-J., 2019. The role of GRHL2 and epigenetic remodeling in epithelial–mesenchymal plasticity in ovarian cancer cells. Commun. Biol. 2, 272. doi:10.1038/s42003-019-0506-3

Cieply, B., Farris, J., Denvir, J., Ford, H.L., Frisch, S.M., 2013. Epithelial-Mesenchymal Transition and Tumor Suppression Are Controlled by a Reciprocal Feedback Loop between ZEB1 and Grainyhead-like-2. Cancer Res. 73, 6299–309. doi:10.1158/0008-5472.CAN-12-4082

Cook, D.P., Vanderhyden, B.C., 2020. Context specificity of the EMT transcriptional response. Nat. Commun. 11, 2142. doi:10.1038/s41467-020-16066-2

Creighton, C.J., Li, X., Landis, M., Dixon, J.M., Neumeister, V.M., Sjolund, A., Rimm, D.L., Wong, H., Rodriguez, A., Herschkowitz, J.I., Fan, C., Zhang, X., He, X., Pavlick, A., Gutierrez, M.C., Renshaw, L., Larionov, A.A., Faratian, D., Hilsenbeck, S.G., Perou, C.M., Lewis, M.T., Rosen, J.M., Chang, J.C., 2009. Residual breast cancers after conventional therapy display mesenchymal as well as tumor-initiating features. Proc. Natl. Acad. Sci. U. S. A. 106, 13820–13825. doi:10.1073/pnas.0905718106

Dave, N., Guaita-Esteruelas, S., Gutarra, S., Frias, A., Beltran, M., Peiro, S., de Herreros, A. G., 2011. Functional cooperation between Snail1 and twist in the regulation of ZEB1 expression during epithelial to mesenchymal transition. J Biol Chem 286, 12024–32. doi:10.1074/jbc.M110.168625

Davis, S., Meltzer, P.S., 2007. GEOquery: A bridge between the Gene Expression Omnibus (GEO) and BioConductor. Bioinformatics 23, 1846–7. doi:10.1093/bioinformatics/btm254

Deshmukh, A.P., Vasaikar, S. V., Tomczak, K., Tripathi, S., Den Hollander, P., Arslan, E., Chakraborty, P., Soundararajan, R., Jolly, M.K., Rai, K., Levine, H., Mani, S.A., 2021. Identification of EMT signaling cross-talk and gene regulatory networks by single-cell RNA sequencing. Proc. Natl. Acad. Sci. U. S. A. 118, e2102050118. doi:10.73/pnas.2102050118

Devaraj, V., Bose, B., 2019. Morphological State Transition Dynamics in EGF-Induced Epithelial to Mesenchymal Transition. J Clin Med 8, 911. doi:10.3390/jcm8070911

Dobin, A., Davis, C.A., Schlesinger, F., Drenkow, J., Zaleski, C., Jha, S., Batut, P., Chaisson, M., Gingeras, T.R., 2013. STAR: Ultrafast universal RNA-seq aligner. Bioinformatics 29, 15–21. doi:10.1093/bioinformatics/bts635

Dong, J., Hu, Y., Fan, X., Wu, X., Mao, Y., Hu, B., Guo, H., Wen, L., Tang, F., 2018. Singlecell RNA-seq analysis unveils a prevalent epithelial/mesenchymal hybrid state during mouse organogenesis. Genome Biol. 19, 31. doi:10.1186/s13059-018-1416-2

Dongre, A., Rashidian, M., Reinhardt, F., Bagnato, A., Keckesova, Z., Ploegh, H.L., Weinberg, R.A., 2017. Epithelial-to-mesenchymal transition contributes to immunosuppression in breast carcinomas. Cancer Res. 77, 3982–3989. doi:10.1158/0008-5472.CAN-16-3292

Drápela, S., Bouchal, J., Jolly, M.K., Culig, Z., 2020. ZEB1 : A Critical Regulator of Cell Plasticity, DNA Damage Response, and Therapy Resistance. Front. Mol. Biosci. 7, 36. doi:10.3389/fmolb.2020.00036

Dumont, N., Wilson, M.B., Crawford, Y.G., Reynolds, P.A., Sigaroudinia, M., Tlsty, T.D., 2008. Sustained induction of epithelial to mesenchymal transition activates DNA methylation of genes silenced in basal-like breast cancers. Proc. Natl. Acad. Sci. 105, 14867–14872. doi:10.1073/pnas.0807146105

Foroutan, M., Bhuva, D.D., Lyu, R., Horan, K., Cursons, J., Davis, M.J., 2018. Single sample scoring of molecular phenotypes. BMC Bioinformatics 19, 404. doi:10.1186/s12859-018-2435-4

Geng, J., Huang, X., Li, Y., Xu, X., Li, S., Jiang, Dingyuan, Liang, J., Jiang, Dianhua, Wang, C., Dai, H., 2015. Down-regulation of USP13 mediates phenotype transformation of fibroblasts in idiopathic pulmonary fibrosis. Respir. Res. 16, 124. doi:10.1186/s12931-015-0286-3

George, J.T., Jolly, M.K., Xu, S., Somarelli, J.A., Levine, H., 2017. Survival outcomes in cancer patients predicted by a partial EMT gene expression scoring metric. Cancer Res. 77, 6415–6428. doi:10.1158/0008-5472.CAN-16-3521

Godin, L., Balsat, C., Eycke, Y. Van, Allard, J., Royer, C., Remmelink, M., Pastushenko, I., Haene, N.D., Blanpain, C., Salmon, I., Rorive, S., Decaestecker, C., 2020. A Novel Approach for Quantifying Cancer Cells Showing Hybrid Epithelial / Mesenchymal States in Large Series of Tissue Samples : Towards a New Prognostic Marker. Cancers (Basel). 12, 906. doi:10.3390/cancers12040906

Gouda, M.M., Shaikh, S.B., Bhandary, Y.P., 2018. Inflammatory and Fibrinolytic System in Acute Respiratory Distress Syndrome. Lung 196, 609–616. doi:10.1007/s00408-018-0150-6

Grande, M.T., Sánchez-Laorden, B., López-Blau, C., De Frutos, C.A., Boutet, A., Arévalo, M., Rowe, R.G., Weiss, S.J., López-Novoa, J.M., Nieto, M.A., 2015. Snail1-induced partial epithelial-to-mesenchymal transition drives renal fibrosis in mice and can be targeted to reverse established disease. Nat. Med. 21, 989–997. doi:10.1038/nm.3901

Guo, C.C., Majewski, T., Zhang, L., Yao, H., Bondaruk, J., Wang, Y., Zhang, S., Wang, Z., Lee, J.G., Lee, S., Cogdell, D., Zhang, M., Wei, P., Grossman, H.B., Kamat, A., Duplisea, J.J., Ferguson, J.E., Huang, H., Dadhania, V., Gao, J., Dinney, C., Weinstein, J.N., Baggerly, K., McConkey, D., Czerniak, B., 2019. Dysregulation of EMT Drives the Progression to Clinically Aggressive Sarcomatoid Bladder Cancer. Cell Rep. 27, 1781–1793.e4. doi:10.1016/j.celrep.2019.04.048

Gupta, G.P., Massague, J., 2006. Cancer Metastasis: Building a Framework. Cell 127, 679–695. doi:10.1016/j.cell.2006.11.001

Hendley, A.M., Rao, A.A., Leonhardt, L., Ashe, S., Smith, J.A., Giacometti, S., Peng, X.L., Jiang, H., Berrios, D.I., Pawlak, M., Li, L.Y., Lee, J., Collisson, E.A., Anderson, M.S., Fragiadakis, G.K., Yeh, J.J., Ye, C.J., Kim, G.E., Weaver, V.M., Hebrok, M., 2021. Single-cell transcriptome analysis defines heterogeneity of the murine pancreatic ductal tree. Elife 10, e67776. doi:10.7554/eLife.67776

Hong, D., Messier, T.L., Tye, C.E., Dobson, J.R., Fritz, A.J., Sikora, K.R., Browne, G., Stein, J.L., Lian, J.B., Stein, G.S., 2017. Runx1 stabilizes the mammary epithelial cell phenotype and prevents epithelial to mesenchymal transition. Oncotarget 8, 17610–17627. doi:10.18632/oncotarget.15381

Huang, R.Y.-J., Wong, M.K., Tan, T.Z., Kuay, K.T., Ng, a H.C., Chung, V.Y., Chu, Y.-S., Matsumura, N., Lai, H.-C., Lee, Y.F., Sim, W.-J., Chai, C., Pietschmann, E., Mori, S., Low, J.J.H., Choolani, M., Thiery, J.P., 2013. An EMT spectrum defines an anoikis-resistant and spheroidogenic intermediate mesenchymal state that is sensitive to e-cadherin restoration by a src-kinase inhibitor, saracatinib (AZD0530). Cell Death Dis. 4, e915. doi:10.1038/cddis.2013.442

Ishay-Ronen, D., Diepenbruck, M., Kalathur, R.K.R., Sugiyama, N., Tiede, S., Ivanek, R., Bantug, G., Morini, M.F., Wang, J., Hess, C., Christofori, G., 2019. Gain Fat—Lose Metastasis: Converting Invasive Breast Cancer Cells into Adipocytes Inhibits Cancer Metastasis. Cancer Cell 35, 17–32.e6. doi:10.1016/j.ccell.2018.12.002

Jia, D., Jolly, M.K., Boareto, M., Parsana, P., Mooney, S.M., Pienta, K.J., Levine, H., Ben-Jacob, E., 2015. OVOL guides the epithelial-hybrid-mesenchymal transition. Oncotarget 6, 15436–48. doi:10.18632/oncotarget.3623

Jia, D., Park, J.H., Kaur, H., Jung, K.H., Yang, S., Tripathi, S., Galbraith, M., Deng, Y., Jolly, M.K., Kaipparettu, B.A., Onuchic, J.N., Levine, H., 2021. Towards decoding the coupled decision-making of metabolism and epithelial-to-mesenchymal transition in cancer. Br. J. Cancer 124, 1902–1911. doi:10.1038/s41416-021-01385-y

Jia, W., Deshmukh, A., Mani, S.A., Jolly, M.K., Levine, H., 2019. A possible role for epigenetic feedback regulation in the dynamics of the Epithelial-Mesenchymal Transition (EMT). Phys. Biol. 16, 066004. doi:10.1088/1478-3975/ab34df

Johnson, K.S., Hussein, S., Chakraborty, P., Muruganantham, A., Mikhail, S., Gonzalez, G., Song, S., Jolly, M.K., Toneff, M.J., Benton, M.L., Lin, Y.C., Taube, J.H., 2021. Epithelial-mesenchymal plasticity through loss of CTCF motif accessibility and protein expression. bioRxiv 447526. doi:10.1101/2021.06.08.447526

Jolly, M., Ward, C., Eapen, M.S., Myers, S., Hallgren, O., Levine, H., Sohal, S.S., 2018. Epithelial-mesenchymal transition, a spectrum of states: Role in lung development, homeostasis, and disease. Dev. Dyn. 247, 346–358. doi:10.1002/dvdy.24541

Jolly, M.K., Boareto, M., Debeb, B.G., Aceto, N., Farach-Carson, M.C., Woodward, W.A., Levine, H., 2017. Inflammatory Breast Cancer: a model for investigating cluster-based dissemination. NPJ Breast Cancer 3, 21. doi:https://doi.org/10.1101/119479

Jolly, M.K., Levine, H., 2017. Computational systems biology of epithelial-hybrid-mesenchymal transitions. Curr. Opin. Syst. Biol. 3, 1–6. doi:10.1016/j.coisb.2017.02.004

Jolly, M.K., Murphy, R., Bhatia, S., Whitfield, H.J., Redfern, A., Davis, M.J., Thompson, E.W., 2021. Measuring and Modelling the Epithelial-Mesenchymal Hybrid State in Cancer: Clinical Implications. Cells Tissues Organs in press. doi:10.1159/000515289

Jolly, M.K., Tripathi, S.C., Jia, D., Mooney, S.M., Celiktas, M., Hanash, S.M., Mani, S.A., Pienta, K.J., Ben-Jacob, E., Levine, H., 2016. Stability of the hybrid epithelial/mesenchymal phentoype. Oncotarget 7, 27067–27084.

Karacosta, L.G., Anchang, B., Ignatiadis, N., Kimmey, S.C., Benson, J.A., Shrager, J.B., Tibshirani, R., Bendall, S.C., Plevritis, S.K., 2019. Mapping Lung Cancer Epithelial-Mesenchymal Transition States and Trajectories with Single-Cell Resolution. Nat. Commun. 10, 5587. doi:10.1101/570341

Kröger, C., Afeyan, A., Mraz, J., Eaton, E.N., Reinhardt, F., Khodor, Y.L., Thiru, P., Bierie, B., Ye, X., Burge, C.B., Weinberg, R.A., 2019. Acquisition of a hybrid E/M state is essential for tumorigenicity of basal breast cancer cells. Proc. Natl. Acad. Sci. 116, 7353–7362. doi:10.1073/pnas.1812876116

Lai, X., Li, Q., Wu, F., Lin, J., Chen, J., Zheng, H., Guo, L., 2020. Epithelial-Mesenchymal Transition and Metabolic Switching in Cancer: Lessons From Somatic Cell Reprogramming. Front. Cell Dev. Biol. 8, 760. doi:10.3389/fcell.2020.00760

Lecharpentier, A., Vielh, P., Perez-Moreno, P., Planchard, D., Soria, J.C., Farace, F., 2011. Detection of circulating tumour cells with a hybrid (epithelial/mesenchymal) phenotype in patients with metastatic non-small cell lung cancer. Br. J. Cancer 105, 1338–1341. doi:10.1038/bjc.2011.405

Leroy, P., Mostov, K.E., 2007. Slug Is Required for Cell Survival during Partial Epithelial-Mesenchymal Transition of HGF-induced tubulogenesis. J. Cell Sci. 18, 1943–1952. doi:10.1091/mbc.E06

Li, H., Courtois, E.T., Sengupta, D., Tan, Y., Chen, K.H., Goh, J.J.L., Kong, S.L., Chua, C., Hon, L.K., Tan, W.S., Wong, M., Choi, P.J., Wee, L.J.K., Hillmer, A.M., Tan, I.B., Robson, P., Prabhakar, S., 2017. Reference component analysis of single-cell transcriptomes elucidates cellular heterogeneity in human colorectal tumors. Nat. Genet. 49, 708–718. doi:10.1038/ng.3818

Li, H., Handsaker, B., Wysoker, A., Fennell, T., Ruan, J., Homer, N., Marth, G., Abecasis, G., Durbin, R., 2009. The Sequence Alignment/Map format and SAMtools. Bioinformatics 25, 2078–2079. doi:10.1093/bioinformatics/btp352

Li, X., Jolly, M.K., George, J.T., Pienta, K.J., Levine, H., 2019. Computational Modeling of the Crosstalk Between Macrophage Polarization and Tumor Cell Plasticity in the Tumor Microenvironment. Front. Oncol. 9, 1–12. doi:10.3389/fonc.2019.00010

Liberzon, A., Subramanian, A., Pinchback, R., Thorvaldsdóttir, H., Tamayo, P., Mesirov, J.P., 2011. Molecular signatures database (MSigDB) 3.0. Bioinformatics 27, 1739–1740. doi:10.1093/bioinformatics/btr260

McFaline-Figueroa, J.L., Hill, A.J., Qiu, X., Jackson, D., Shendure, J., Trapnell, C., 2019. A pooled single-cell genetic screen identifies regulatory checkpoints in the continuum of the epithelial-to-mesenchymal transition. Nat. Genet. 51, 1389–1398. doi:10.1038/s41588-019-0489-5

Mooney, S.M., Talebian, V., Jolly, M.K., Jia, D., Gromala, M., Levine, H., McConkey, B.J., 2017. The GRHL2/ZEB Feedback Loop—A Key Axis in the Regulation of EMT in Breast Cancer. J. Cell. Biochem. 118, 2559–2570. doi:10.1002/jcb.25974

Morel, A.-P., Lièvre, M., Thomas, C., Hinkal, G., Ansieau, S., Puisieux, A., 2008. Generation of breast cancer stem cells through epithelial-mesenchymal transition. PLoS One 3, e2888. doi:10.1371/journal.pone.0002888

Neelakantan, D., Zhou, H., Oliphant, M.U.J., Zhang, X., Simon, L.M., Henke, D.M., Shaw, C. A., Wu, M.F., Hilsenbeck, S.G., White, L.D., Lewis, M.T., Ford, H.L., 2017. EMT cells increase breast cancer metastasis via paracrine GLI activation in neighbouring tumour cells. Nat. Commun. 8, 15773. doi:10.1038/ncomms15773

Nieto, M.A., Huang, R.Y., Jackson, R.A., Thiery, J.P., 2016. EMT: 2016. Cell 166, 21–45. doi:10.1016/j.cell.2016.06.028

Pal, A., Barrett, T.F., Paolini, R., Parikh, A., Puram, S. V., 2021. Partial EMT in head and neck cancer biology: a spectrum instead of a switch. Oncogene 40, 5049–5065. doi:10.1038/s41388-021-01868-5

Pasani, S., Sahoo, S., Jolly, M.K., 2021. Hybrid E/M phenotype(s) and stemness: a mechanistic connection embedded in network topology. J Clin Med 10, 60. doi:10.1101/2020.10.18.341271

Pastushenko, I., Blanpain, C., 2019. EMT Transition States during Tumor Progression and Metastasis. Trends Cell Biol. 29, 212–226. doi:10.1016/j.tcb.2018.12.001

Pastushenko, I., Brisebarre, A., Sifrim, A., Fioramonti, M., Revenco, T., Boumahdi, S., Van Keymeulen, A., Brown, D., Moers, V., Lemaire, S., De Clercq, S., Minguijón, E., Balsat, C., Sokolow, Y., Dubois, C., De Cock, F., Scozzaro, S., Sopena, F., Lanas, A., D’Haene, N., Salmon, I., Marine, J.-C., Voet, T., Sotiropoulou, P.A., Blanpain, C., 2018. Identification of the tumour transition states occurring during EMT. Nature 556, 463–468. doi:10.1038/s41586-018-0040-3

Peixoto, P., Etcheverry, A., Aubry, M., Missey, A., Lachat, C., Perrard, J., Hendrick, E., Delage-Mourroux, R., Mosser, J., Borg, C., Feugeas, J.P., Herfs, M., Boyer-Guittaut, M., Hervouet, E., 2019. EMT is associated with an epigenetic signature of ECM remodeling genes. Cell Death Dis. 10. doi:10.1038/s41419-019-1397-4

Puram, S. V., Tirosh, I., Parikh, A.S., Patel, A.P., Yizhak, K., Gillespie, S., Rodman, C., Luo, C.L., Mroz, E.A., Emerick, K.S., Deschler, D.G., Varvares, M.A., Mylvaganam, R., Rozenblatt-Rosen, O., Rocco, J.W., Faquin, W.C., Lin, D.T., Regev, A., Bernstein, B.E., 2017. Single-Cell Transcriptomic Analysis of Primary and Metastatic Tumor Ecosystems in Head and Neck Cancer. Cell 171, 1611–1624. doi:10.1016/j.cell.2017.10.044

Ruscetti, M., Dadashian, E.L., Guo, W., Quach, B., Mulholland, D.J., Park, J.W., Tran, L.M., Kobayashi, N., Bianchi-Frias, D., Xing, Y., Nelson, P.S., Wu, H., 2016. HDAC inhibition impedes epithelial-mesenchymal plasticity and suppresses metastatic, castrationresistant prostate cancer. Oncogene 35, 3781–95. doi:10.1038/onc.2015.444

Sahoo, S., Mishra, A., Kaur, H., Hari, K., Muralidharan, S., Mandal, S., Jolly, M.K., 2021a. A mechanistic model captures the emergence and implications of non-genetic heterogeneity and reversible drug resistance in ER+ breast cancer cells. NAR Cancer 3. doi:10.1093/narcan/zcab027

Sahoo, S., Nayak, S.P., Hari, K., Purkait, P., Mandal, S., Kishore, A., Levine, H., Jolly, M.K., 2021b. Immunosuppressive traits of the hybrid epithelial/mesenchymal phenotype. bioRxiv 449285. doi:10.1101/2021.06.21.449285

Saxena, K., Subbalakshmi, A.R., Jolly, M.K., 2019. Phenotypic heterogeneity in circulating tumor cells and its prognostic value in metastasis and overall survival. EBioMedicine 46, 4–5. doi:10.1016/j.ebiom.2019.07.074

Serresi, M., Kertalli, S., Li, L., Schmitt, M.J., Dramaretska, Y., Wierikx, J., Hulsman, D., Gargiulo, G., 2021. Functional antagonism of chromatin modulators regulates epithelial-mesenchymal transition. Sci. Adv. 7, eabd7974. doi:10.1126/sciadv.abd7974

Simeonov, K.P., Byrns, C.N., Clark, M.L., Norgard, R.J., Martin, B., Stanger, B.Z., Shendure, J., McKenna, A., Lengner, C.J., 2021. Single-cell lineage tracing of metastatic cancer reveals selection of hybrid EMT states. Cancer Cell 39, 1150–1162.e9. doi:10.1016/j.ccell.2021.05.005

Sivakumar, P., Thompson, J.R., Ammar, R., Porteous, M., McCoubrey, C., Cantu, E., Ravi, K., Zhang, Y., Luo, Y., Streltsov, D., Beers, M.F., Jarai, G., Christie, J.D., 2019. RNA sequencing of transplant-stage idiopathic pulmonary fibrosis lung reveals unique pathway regulation. ERJ Open Res. 5, 00117–02019. doi:10.1183/23120541.00117-2019

Sohal, S.S., 2017. Epithelial and endothelial cell plasticity in chronic obstructive pulmonary disease (COPD). Respir. Investig. 55, 104–113. doi:10.1016/j.resinv.2016.11.006

Stylianou, N., Lehman, M.L., Wang, C., Fard, A.T., Rockstroh, A., Fazli, L., Jovanovic, L., Ward, M., Sadowski, M.C., Kashyap, A.S., Buttyan, R., Gleave, M.E., Westbrook, T.F., Williams, E.D., Gunter, J.H., Nelson, C.C., Hollier, B.G., 2019. A molecular portrait of epithelial–mesenchymal plasticity in prostate cancer associated with clinical outcome. Oncogene 38, 913–934. doi:10.1038/s41388-018-0488-5

Subramanian, A., Tamayo, P., Mootha, V.K., Mukherjee, S., Ebert, B.L., Gillette, M.A., Paulovich, A., Pomeroy, S.L., Golub, T.R., Lander, E.S., Mesirov, J.P., 2005. Gene set enrichment analysis: A knowledge-based approach for interpreting genome-wide expression profiles. Proc Natl Acad Sci U S A 102, 15545–15550. doi:10.1073/pnas.0506580102

Tan, T.Z., Miow, Q.H., Miki, Y., Noda, T., Mori, S., Huang, R.Y., Thiery, J.P., 2014. Epithelial-mesenchymal transition spectrum quantification and its efficacy in deciphering survival and drug responses of cancer patients. EMBO Mol. Med. 6, 1279–1293. doi:10.15252/emmm.201404208

Tian, B., Li, X., Kalita, M., Widen, S.G., Yang, J., Bhavnani, S.K., Dang, B., Kudlicki, A., Sinha, M., Kong, F., G, W.T., Luxon, B.A., Brasier, A.R., 2015. Analysis of the TGFβ-induced program in primary airway epithelial cells shows essential role of NF-κB/RelA signaling network in type II epithelial mesenchymal transition. BMC Genomics 16, 529. doi:10.1186/s12864-015-1707-x

Tripathi, S.C., Peters, H.L., Taguchi, A., Katayama, H., Wang, H., Momin, A., Jolly, M.K., Celiktas, M., Rodriguez-Canales, J., Liu, H., Behrens, C., Wistuba, I.I., Ben-Jacob, E., Levine, H., Molldrem, J.J., Hanash, S.M., Ostrin, E.J., 2016. Immunoproteasome deficiency is a feature of non-small cell lung cancer with a mesenchymal phenotype and is associated with a poor outcome. Proc. Natl. Acad. Sci. 113, E1555–64. doi:10.1073/pnas.1521812113

Tripathi, V., Sixt, K.M., Gao, S., Xu, X., Huang, J., Weigert, R., Zhou, M., Zhang, Y.E., 2016. Direct Regulation of Alternative Splicing by SMAD3 through PCBP1 Is Essential to the Tumor-Promoting Role of TGF-β. Mol. Cell 64, 549–564. doi:10.1016/j.molcel.2016.09.013

Tsuji, T., Ibaragi, S., Shima, K., Hu, M.G., Katsurano, M., Sasaki, A., Hu, G.F., 2008. Epithelial-mesenchymal transition induced by growth suppressor p12 CDK2-AP1 promotes tumor cell local invasion but suppresses distant colony growth. Cancer Res. 68, 10377–10386. doi:10.1158/0008-5472.CAN-08-1444

Vasaikar, S. V., Deshmukh, A.P., den Hollander, P., Addanki, S., Kuburich, N.A., Kudaravalli, S., Joseph, R., Chang, J.T., Soundararajan, R., Mani, S.A., 2021. EMTome: a resource for pan-cancer analysis of epithelial-mesenchymal transition genes and signatures. Br. J. Cancer 124, 259–269. doi:10.1038/s41416-020-01178-9

Wang, P., Zhou, Renwu, Thomas, P., Zhao, L., Zhou, Rusen, Mandal, S., Jolly, M.K., Richard, D.J., Rehm, B.H.A., Ostrikov, K., Dai, X., Williams, E.D., Thompson, E.W., 2021. Epithelial-to-mesenchymal transition enhances cancer cell sensitivity to cytotoxic effects of zcold atmospheric plasmas in breast and bladder cancer systems. Cancers (Basel). 13, 2889. doi:10.3390/cancers13122889

Wang, W., Douglas, D., Zhang, J., Kumari, S., Enuameh, M.S., Dai, Y., Wallace, C.T., Watkins, S.C., Shu, W., Xing, J., 2020. Live-cell imaging and analysis reveal cell phenotypic transition dynamics inherently missing in snapshot data. Sci. Adv. 6, eaba9319. doi:10.1126/sciadv.aba9319

Wang, Z., Li, Y., Kong, D., Banerjee, S., Ahmad, A., Azmi, A.S., Ali, S., Abbruzzese, J.L., Gallick, G.E., Sarkar, F.H., 2009. Acquisition of epithelial-mesenchymal transition phenotype of gemcitabine-resistant pancreatic cancer cells is linked with activation of the notch signaling pathway. Cancer Res. 69, 2400–7. doi:10.1158/0008-5472.CAN-08-4312

Watanabe, K., Panchy, N., Noguchi, S., Suzuki, H., Hong, T., 2019. Combinatorial perturbation analysis reveals divergent regulations of mesenchymal genes during epithelial-to-mesenchymal transition. npj Syst. Biol. Appl. 5, 21. doi:10.1038/s41540-019-0097-0

Yang, H., Adam, R.C., Ge, Y., Hua, Z.L., Fuchs, E., 2017. Epithelial-Mesenchymal Microniches Govern Stem Cell Lineage Choices. Cell 169, 483–496.e13. doi:10.1016/j.cell.2017.03.038

Yang, J., Mani, S.A., Donaher, J.L., Ramaswamy, S., Itzykson, R.A., Come, C., Savagner, P., Gitelman, I., Richardson, A., Weinberg, R.A., Val, C., Lamarque, A., 2004. Twist, a master regulator of Morphogenesis, plays an essential role in tumor metastasis. Cell 117, 927–939. doi:10.1016/j.cell.2004.06.006

Yu, M., Bardia, A., Wittner, B.S., Stott, S.L., Smas, M.E., Ting, D.T., Isakoff, S.J., Ciciliano, J.C., Wells, M.N., Shah, A.M., Concannon, K.F., Donaldson, M.C., Sequist, L. V, Brachtel, E., Sgroi, D., Baselga, J., Ramaswamy, S., Toner, M., Haber, D.A., Maheswaran, S., 2013. Circulating breast tumor cells exhibit dynamic changes in epithelial and mesenchymal composition. Science 339, 580–4. doi:10.1126/science.1228522

Zhang, S., Cui, Y., Ma, X., Yong, J., Yan, L., Yang, M., Ren, J., Tang, F., Wen, L., Qiao, J., 2020. Single-cell transcriptomics identifies divergent developmental lineage trajectories during human pituitary development. Nat. Commun. 11, 5275. doi:10.1038/s41467-020-19012-4

Zhao, S., Fung-Leung, W.P., Bittner, A., Ngo, K., Liu, X., 2014. Comparison of RNA-Seq and microarray in transcriptome profiling of activated T cells. PLoS One 9, e78644. doi:10.1371/journal.pone.0078644

